# Improved Super-Resolution Ribosome Profiling Revealed Prevalent Translation of Upstream ORFs and Small ORFs in Arabidopsis

**DOI:** 10.1101/2023.09.08.556947

**Authors:** Hsin-Yen Larry Wu, Qiaoyun Ai, Rita Teresa Teixeira, Gaoyuan Song, Christian Montes, J. Mitch Elmore, Justin W. Walley, Polly Yingshan Hsu

## Abstract

A crucial step in functional genomics is identifying actively translated open reading frames (ORFs) that link biological functions. The challenge lies in identifying short ORFs, as they are greatly impacted by data quality and depth. Here, we improved the coverage of super-resolution Ribo-seq in Arabidopsis, revealing uncharacterized translation events in nucleus-, chloroplast-, and mitochondria-encoded genes. We identified 7,751 unconventional translation events, including 6,996 upstream ORFs (uORFs) and 209 downstream ORFs on annotated protein-coding genes, as well as 546 ncORFs on presumed non-coding RNAs. Proteomics data confirmed the production of stable proteins from some of the unannotated translation events. We present evidence of active translation on primary transcripts of tasiRNAs (*TAS1-4*) and microRNAs (pri-miR163, pri-miR169), and periodic ribosome stalling supporting co-translational decay. Additionally, we developed a method for identifying extremely short uORFs, including 370 minimum uORF (AUG-stop), and 2,984 tiny uORFs (2-10 aa), as well as 681 uORFs that overlap with each other. Remarkably, these short uORFs exhibit strong translational repression as longer uORFs. We also systematically discovered 594 uORFs regulated by alternative splicing, suggesting widespread isoform-specific translational control. Finally, these prevalent uORFs are associated with numerous important pathways. In summary, our improved Arabidopsis translational landscape provides valuable resources to study gene expression regulation.

## INTRODUCTION

Accurately defining gene models and determining translated open reading frames (ORFs) is fundamental for studying gene functions and monitoring cellular activity in all living organisms. Despite extensive efforts, our understanding of the translational landscape remains incomplete (Andrews and Rothnagel, 2014; Hellens et al., 2016; Orr et al., 2020; Wu et al., 2023). It has become increasingly clear that a substantial fraction of the transcriptomes in diverse plant species has prevalent unannotated translated ORFs. These unannotated translated ORFs include small ORFs (sORFs) encoded by presumed non-coding RNAs, as well as short upstream ORFs (uORFs) present in the 5′ untranslated regions (5′ UTRs) of protein-coding RNAs (i.e., mRNAs) (Liu et al., 2013; Juntawong et al., 2014; Lei et al., 2015; Hsu et al., 2016; Bazin et al., 2017; Wu et al., 2019; Li and Liu, 2020; Kurihara et al., 2020; Sotta et al., 2022; Guo et al., 2023; Qanmber et al., 2023; Zhu et al., 2023). More recently, even downstream ORFs (dORFs) in the 3′ UTRs of mRNAs have been reported in vertebrates (Wu et al., 2020). Serendipitous discoveries from early forward genetics and recent experimental evidence have shown that these relatively short ORFs can produce small proteins or peptides that play important roles in various aspects of signaling and physiology (Tavormina et al., 2015; Hsu and Benfey, 2018; Takahashi et al., 2019). In addition, the act of translation itself may have regulatory roles, even if the protein products are not functional (Orr et al., 2020). For example, uORFs have long been recognized as cis-regulatory elements suppressing protein synthesis of the downstream main ORFs (mORFs) and are associated with various plant phenotypes (Von Arnim et al., 2014; Xu et al., 2017; Zhang et al., 2018; Xing et al., 2020; Gage et al., 2022). More recently, the ribosome association with sORFs on the primary transcripts of trans-acting small interfering RNAs (tasiRNAs) has been suggested to bring the transcripts to the rough endoplasmic reticulum (ER) for microRNA (miRNA)-mediated tasiRNAs biogenesis (Li et al., 2016; Yoshikawa et al., 2016; Hou et al., 2016; Bazin et al., 2017; Iwakawa et al., 2021).

Despite their importance, short ORFs are commonly excluded by computational genome annotations. As a vast number of potential ORF sequences can occur randomly, computational annotations rely on several assumptions about ORFs to minimize false positive identifications. These assumptions typically include the translation of only one ORF, the longest one, from a single transcript, and the requirement for the ORF to be greater than 100 codons (Basrai et al., 1997; Olsen et al., 2002; Lease and Walker, 2006). Although this approach is robust and efficient, genuine mRNAs encoding sORFs can be misannotated as non-coding RNAs. Moreover, the polycistronic potential of mRNAs is not considered, resulting in the exclusion of uORFs or dORFs in the annotation. Thus, comprehensive identification of these hidden short ORFs is the first step toward understanding and characterizing their functions.

Bioinformatic approaches based on evolutionary conservation or sequence homology to known sORFs have been attempted (Olsen et al., 2002; Lease and Walker, 2006; Hanada et al., 2007, 2009; Zhou et al., 2013; de Bang et al., 2017; Feng et al., 2023; Li et al., 2023). However, accumulating evidence suggests that many sORFs have evolved recently during evolution (Ruiz-Orera et al., 2014; Sandmann et al., 2023), and some sORFs are only conserved in specific families or small groups of plants. For instance, the first plant peptide hormone involved in the wounding response in tomato, systemin, is only present in Solaneae, a subtribe of the Solanaceae (Pearce et al., 1991; Constabel et al., 1998). Additionally, a small protein called Qua Quine Starch (QQS, 59 aa), involved in carbon and nitrogen allocation as well as pathogen susceptibility across species, is encoded by an orphan gene that exists only in Arabidopsis (Li et al., 2009, 2015; Qi et al., 2019). In our previous analysis of the tomato translatome, we found many translated sORFs specific to the Solanaceae or exclusively found in tomato (Wu et al., 2019).

Similarly, although 30-70% of plant genes contain potential uORFs, only 119 uORFs encoding <u>c</u>onserved <u>p</u>eptides (CPuORFs) corresponding to 81 homology groups, have been identified in Arabidopsis despite extensive searches (Hayden and Jorgensen, 2007; Jorgensen and Dorantes-Acosta, 2012; Vaughn et al., 2012; Takahashi et al., 2012; Van Der Horst et al., 2019, 2020; Takahashi et al., 2020; Zhang et al., 2021). Some CPuORFs are also specific to certain plant families. For example, *MYB51* contains a Brassicaceae-specific CPuORF. This is likely due to the fact that MYB51 regulates the biosynthesis of glucosinolates, a secondary metabolite mainly found in the order of Brassicales (Hou et al., 2016). Therefore, these family-or species-specific sORFs and uORFs may have evolved to provide functions specific to certain groups of plants. Together, the small size, low conservation, and a limited number of experimentally characterized sORFs and uORFs available as training datasets restrict the power of bioinformatic prediction. For these reasons, a systematic experimental approach is necessary to uncover these hidden functional short ORFs.

Ribosome profiling (Ribo-seq) has emerged as a high-throughput approach for ORF discovery and quantification. Essentially, Ribo-seq combines ribosome footprinting with deep sequencing (Ingolia et al., 2009; Brar and Weissman, 2015). The procedure involves treating ribosome-bound mRNAs with ribonuclease to obtain ribosome-protected mRNA fragments (RPFs), or ribosome footprints. Sequencing RPFs enables the identification and quantification of ribosome occupancy on mRNAs throughout the transcriptome. Importantly, when using one nucleotide (nt) to assign the position of RPFs on mRNAs, precisely digested RPFs exhibit enrichment in the expected reading frame along the coding sequences, which is called 3-nt periodicity. This periodic property reflects that ribosomes decode 3 nt at a time and is a benchmark for high-quality Ribo-seq data (Ingolia et al., 2009; Jiang et al., 2022). 3-nt periodicity has been considered a reliable feature to distinguish real RPFs from contaminant RNA fragments protected by non-ribosomal protein complexes and to separate actively translating ribosomes from ribosomes stalled at certain regions of transcripts without engaging in translation (Guttman et al., 2013; Guydosh and Green, 2014; Jiang et al., 2022). For these reasons, the majority of ORF discovery software utilizes 3-nt periodicity as a key parameter to identify translated ORFs (Wang et al., 2019). Notably, imprecise digestion of RPFs can lead to out-of-frame mapping, reducing the resolution of Ribo-seq data and the confidence of active translation. Therefore, strong 3-nt periodicity is critical for the success of ORF identification.

We have previously optimized the footprinting buffer to improve the precise digestion of RPFs, resulting in over 90% of RPFs enriched in the expected reading frame in Arabidopsis seedling roots and shoots (Hsu et al., 2016). Although the high-quality data revealed numerous unannotated translation events, the number of relatively short ORFs, such as sORFs and uORFs, identified based on significant 3-nt periodicity (Calviello et al., 2016), was low (32 sORFs and 187 uORFs) (Hsu et al., 2016). This is presumably due to insufficient coverage within these short ORFs. Additionally, multiple high-quality Ribo-seq datasets, including those in zebrafish, Chlamydomonas, Arabidopsis, and tomato (Bazzini et al., 2014; Chung et al., 2015; Hsu et al., 2016; Wu et al., 2019), have revealed specific out-of-frame mapping of RPFs at translation termination, likely resulted from structural rearrangement upon binding of eukaryotic Release Factors (eRFs) (Alkalaeva et al., 2006; Brown et al., 2015; Matheisl et al., 2015). This out-of-frame mapping at termination can significantly reduce the overall 3-nt periodicity of short ORFs. Furthermore, identifying uORFs is considered more challenging as they can be extremely short and often overlap with other uORFs, leading to low 3-nt periodicity. Therefore, it was proposed that more specialized software is needed for uORF identification (Wang et al., 2020).

Here, we present an improved Ribo-seq dataset with enhanced RPF coverage in Arabidopsis seedlings. Our new data significantly improves the identification of translated uORFs and sORFs, and provides evidence for supporting noncanonical translation associated with various gene regulation events. Additionally, we developed a new computational method to address the issues of out-of-frame mapping at translation termination and overlapping between uORFs for uncovering relatively short uORFs. Combining our new data and new computational approach has identified over 7000 unannotated translation events. Thus, our enhanced Arabidopsis translational landscape facilitates the discovery of translated ORFs and offers valuable resources for studying gene functions.

## RESULTS

### Enhanced super-resolution Ribo-seq improved the read coverage

Although our previous super-resolution Ribo-seq data revealed many unannotated translation events, only 187 uORFs and 32 sORFs were detected based on significant 3-nt periodicity (Hsu et al., 2016), likely due to insufficient RPF coverage within these short ORFs. We proposed that increasing RPF coverage could improve the efficiency of identifying translated ORFs, especially for relatively short ORFs. We tested a new protocol using 7-day-old Arabidopsis seedlings. Both Ribo-seq and RNA-seq libraries were generated. We made two major modifications in the Ribo-seq library preparation, including 1) changing the order between RPF size selection and rRNA depletion to maximize the input materials, and 2) reducing the steps of RNA purification to minimize potential loss of RPFs (see MATERIALS AND METHODS for details). We observed that the number of PCR cycles needed for the final Ribo-seq library amplification reduced from 12 cycles (Hsu et al., 2016) to 9 cycles, suggesting the RPF yield was increased.

After sequencing and data analysis, we observed excellent correlations among biological replicates for both Ribo-seq and RNA-seq samples, respectively (**Figure S1**, r = 0.99–1 for Ribo-seq; r = 0.98–1 for RNA-seq). Overall, the Ribo-seq samples show good correlations with the RNA-seq samples (**Figure S1**, r = 0.88–0.93), as previously observed (Hsu et al., 2016).

Importantly, our new Ribo-seq data display characteristics expected for high-quality data, including strong 3-nt periodicity (92% in-frame reads for 28-nt RPFs), high enrichment for coding sequences (CDSs), and expected RPF lengths with 28 nt being the most abundant (**Figure 1A–C**). To examine RPF diversity and coverage globally, we next examined how many P-sites (the peptidyl tRNA-binding sites within translating ribosomes) these RPFs mapped to. From 20 million randomly selected reads, the number of unique P-sites detected in our new data is 4.01-fold higher than that in our previous dataset in Arabidopsis seedling shoots and roots grown under similar conditions (**Figure 1D**), indicating an increased RPF diversity in the new data. The coverage improvement is also evident in individual transcript profiles when comparing genes with similar mRNA levels. The overall RPF coverage within mORFs (and expected uORFs, if any) is improved, and RPF levels are less noisy in our current data compared to the previous data (**Figure 1D**, and **8A–B**).

**Figure 1.**
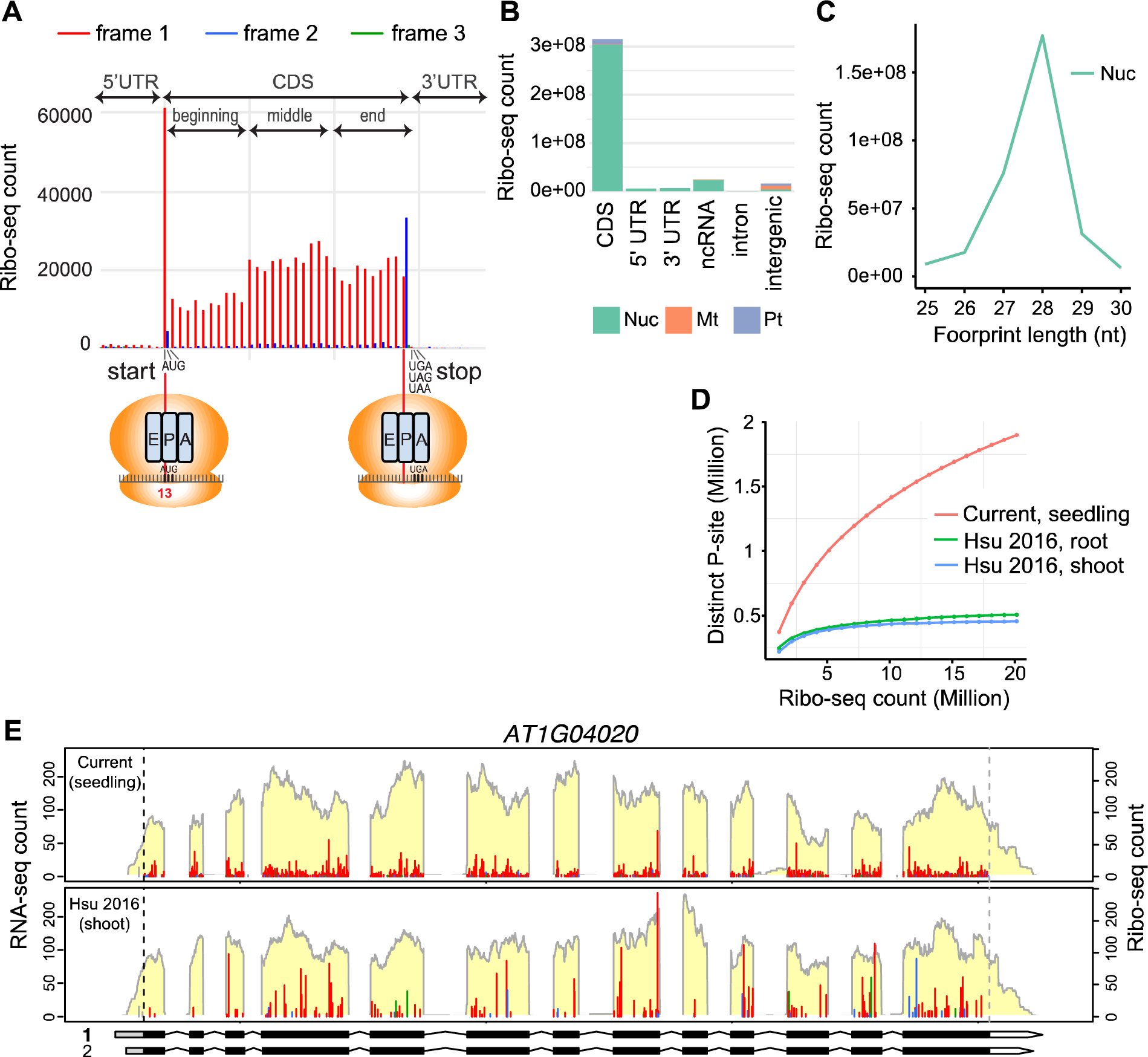
Enhanced Ribo-seq data with improved coverage. (A) Metagene analysis of 28-nt RPFs mapped to regions near the start and stop codons of annotated ORFs in Araport 11. The RPFs are presented with their first nt at the P-site, which is the 13th nt for 28-nt RPFs. The RPFs are colored in red, blue, and green to indicate they are in the first (expected), second, and third reading frames, respectively. The majority of footprints were mapped to the CDS in the expected reading frame (92% in frame). (B) Genomic features mapped by RPFs. Reads that mapped to nuclear (Nuc)-, mitochondrial (Mt)-, and plastid (Pt)-encoded genes are shown. (C) Length distribution of RPFs. Reads that mapped to nuclear-encoded genes are presented. (D) Distinct P-sites detected in 1 to 20 million randomly selected RPFs from our current and previous datasets (Hsu et al., 2016). (E) RNA-seq and Ribo-seq profiles of *AT1G04020* from the current study and our previous shoot data in (Hsu et al., 2016) are presented. RNA-seq coverage is shown with a light-yellow background. Ribo-seq reads are presented with their first nt at the P-site, and they are colored in red, blue, and green to indicate they are in the first (expected), second, and third reading frames, respectively. Reads outside of the ORF range are colored in grey. Within the gene models, black boxes represent the annotated mORFs, and gray and white regions indicate 5′ UTRs and 3′ UTRs, respectively. The specific isoform being plotted is indicated to the left of the gene model and bolded. Black and gray vertical dashed lines represent the translation start and stop, respectively, for the annotated mORF.

### ORF identification based on significant 3-nt periodicity

As the six samples are highly correlated (**Figure S1**), we pooled them to create one large Ribo-seq dataset and one large RNA-seq dataset for ORF identification, with the goal of identifying short ORFs and ORFs potentially translated at lower levels. In total, we have 298 million mapped Ribo-seq reads, corresponding to 11.2 million unique P-sites.

To capture translation events arising from unannotated transcripts, we also performed a reference-guided *de novo* transcriptome assembly using Araport11 annotation (**Figure S2**, workflow for ORF identification; **File S1**, newly assembled transcriptome GTF). We then used RiboTaper (Calviello et al., 2016), a spectrum analysis software, to assess whether a significant 3-nt periodicity is detected within each potential AUG-initiated ORF along both the transcripts annotated in Araport11 and the newly assembled ones.

Our new data dramatically increase the number of translated ORFs detected by RiboTaper (using the default cutoff, Multitaper F-test, p < 0.05). Among annotated protein-coding genes, we identified the translation of 2113 uORFs, 35191 annotated ORFs (referred to as ‘Conventional CDSs (CCDSs)’ by RiboTaper), and 209 dORFs (**Figure 2A**, 2B and **Dataset S1A–C**). Additionally, we identified 546 ORFs (referred to as ‘ncORFs’ by RiboTaper) from transcripts that were not previously considered as coding (**Figure 2A**, **2B** and **Dataset S1D**).

**Figure 2.**
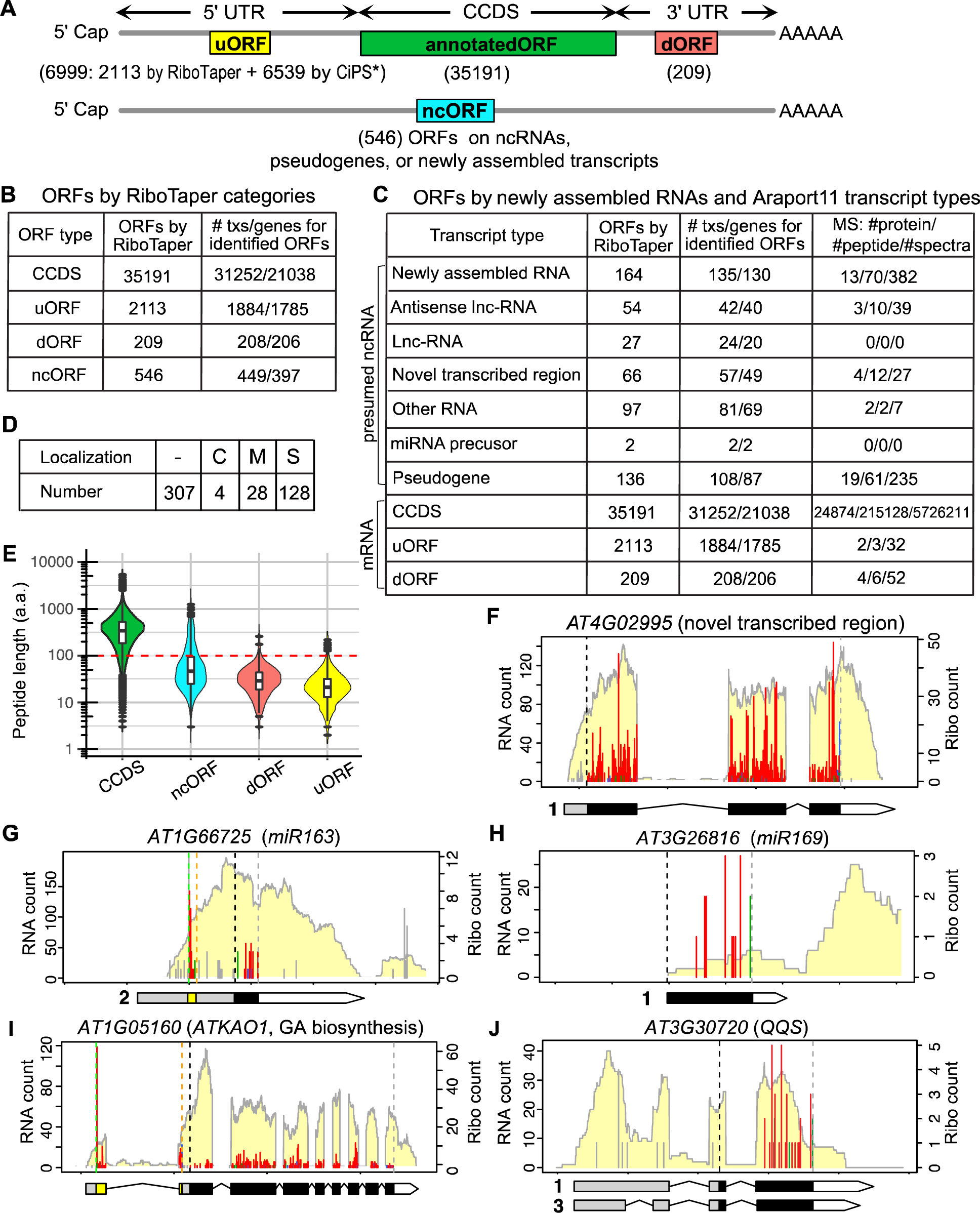
Translated ORFs identified in this study. (A) The number and position of ORFs detected within annotated protein-coding mRNAs and RNAs presumed to be non-coding. Most ORFs were identified by RiboTaper. * Additional uORFs were identified by a separate method, CiPS (see below). (B) The categories of translated ORFs identified by RiboTaper. RiboTaper defines ncORFs as ORFs detected within presumed non-coding RNAs. (C) The number of translated ORFs and proteins identified from either newly assembled RNAs or different annotated transcript types. The 4^th^ column indicates the number of proteins, peptides, and spectrums detected by mass spectrometry for each class of ORFs. Txs: transcripts; MS: mass spectrometry. (D) Subcellular localization of ncORFs of length between 20 and 100 amino acids predicted by TargetP. C: chloroplast, M: mitochondria, S: secreted. (E) Size distribution of annotated ORFs (CCDSs) and other ORFs identified by RiboTaper. (F–J) Examples of translated ORFs in various transcript types. Ribo-seq and RNA-seq profiles are presented in the same way as described in Figure 1E. Additional ORF in (G) or uORF in (I) are shown by a yellow box in the gene model, and their translation start and stop are indicated by green and orange vertical dashed lines, respectively, within the profiles. (F) A translated ORF in a novel transcribed region defined by Araport 11. (G–H) Translated ORFs in primary transcripts of miR163 and miR169. The first ORF within miR163 was visually identified. (I) A translated uORF in *ATKAO1*, which is involved in GA biosynthesis. (J) A translated ORF in *QQS*, a sORF identified from an orphan gene in a previous study (Li et al., 2009).

These ncORFs include 164 from newly assembled transcripts, 246 from annotated lncRNAs, miRNA precursors, other RNAs, etc., and 136 from annotated pseudogenes (**Figure 2C**). The number of ORFs identified in this study is in strong contrast to our previous datasets (187 uORFs, 10 dORFs, 64 ncORFs, etc.). Overall, these unannotated ORFs tend to be small in size. The medians of encoded peptides from ncORFs, dORFs, and uORFs are 46, 29, and 21 amino acids (aa), respectively (**Figure 2E**), which is consistent with the notion that ORFs smaller than 100 aa are often excluded from the annotation (Basrai et al., 1997).

Proteomic analysis using mass spectrometry confirmed the production of stable proteins from at least some of these ncORFs, dORFs, and sORFs, as evidenced by the detection of multiple peptides from each ORF (4^th^ column in **Figure 2C** and **Dataset S2A–B**). Among the 467 ncORFs that are at least 20 aa, TargetP v2.0 (Armenteros et al., 2019) predicted that 4, 28 and 128 of them are targeted to chloroplasts, mitochondria, or are secreted, respectively (**Figure 2D, Dataset S3**), suggesting their potential function in these subcellular localizations.

The evolutionary conservation of 255 single-exon ncORFs was analyzed using tBLASTn. Notably, only 22 of them have homologs outside of the Brassicaceae among the 13 genomes we searched (**Figure S3**). These results are consistent with our previous findings in tomato, where most of the identified tomato sORFs are family or species-specific (Wu et al., 2019). Together these observations align with the notions that many sORFs are likely *de novo* genes that have emerged recently in evolution in other eukaryotes (Ruiz-Orera et al., 2014; Sandmann et al., 2023).

We investigated these unannotated ORFs individually using RiboPlotR (Wu and Hsu, 2021) to evaluate their translation patterns. RiboPlotR displays Ribo-seq and RNA-seq data in the context of gene and transcript structures, and the RPFs within ORFs are color-coded according to the reading frame. Examples of ORFs translated from the Araport11 annotated novel transcribed regions (**Figure 2F**), the primary transcripts of miR163 and miR169 (**Figure 2G–H**), and a newly identified uORF (**Figure 2I**) were presented. We also confirmed the translation of *QQS* (**Figure 2J**), a functional sORF identified in a previous study (Li et al., 2009). It has been shown that the application of synthetic peptides encoded by the sORFs within the primary transcripts of miRNAs (including miR169) increases the expression of the corresponding miRNAs (Lauressergues et al., 2015). However, to our knowledge, no significant RPFs or 3-nt periodicity had been detected in the primary transcripts of miRNAs in previous Ribo-seq data in plants. The two ORFs detected in pri-miR163 encode 9 and 25 aa, respectively (**Figure 2G**), while the ORF detected in pri-miR169 encodes 50 aa (**Figure 2H**). Although these three peptides are not conserved, the second peptide from pri-miR163 is predicted to target to mitochondria according to TargetP.

Taken together, our new data with improved coverage have expanded the number of identified translated ORFs, particularly those of smaller size that are often overlooked in the annotation.

### Translation detected in plastids and mitochondria

In addition to RPFs mapped to nucleus-encoded genes (hereafter referred to as ‘nucleus RPFs’), the sample preparation of our current study also included chloramphenicol, enabling us to examine the RPFs mapped to plastid/chloroplast- and mitochondria-encoded genes (hereafter referred to as ‘plastid RPFs’ and ‘mitochondria RPFs’). Consistent with previous observations (Chotewutmontri and Barkan, 2016), plastid RPFs and mitochondria RPFs only compose a small fraction of total RPFs (**Figure 1B** and **S4**).

Unlike the nucleus RPFs, which are mainly mapped to CDS (**Figure 1B** and **S4**), Ribo-seQC analysis (Calviello et al., 2019) revealed that a substantial fraction of plastid and mitochondria RPFs are mapped to intergenic regions, particularly 33-nt for plastid RPFs and 28-nt for mitochondria RPFs (**Figure S4**). For RPFs mapped to CDS, the predominant RPF length is 28-nt for nucleus and chloroplast RPFs, while it is 29-nt for mitochondria RPFs (**Figure S4**). Overall, the size distribution of these organelle RPFs is similar to previous observations reported in maize and Arabidopsis for plastid and mitochondrial RPFs (Chotewutmontri and Barkan, 2016; Planchard et al., 2018).

We analyzed the frame enrichment in individual RPF lengths using Ribo-seQC and found that 28 nt provides the highest frame enrichment in all three genomes (**Figure S5**). However, at this length, nucleus and plastid RPFs use frame 1 (red), whereas mitochondria RPFs use frame 3 (green) (**Figure S5**). In general, plastid RPFs exhibit strong 3-nt periodicity across the 22-30 nt range, with 71-83% RPFs in frame. In contrast, mitochondria RPFs display relatively weak 3-nt periodicity, with the best in-frame percentage being around 52% at either 28 nt or 22 nt (**Figure S5**).

We also examined the RPFs mapped to annotated start codons to infer the P-site position in each RPF length. Overall, the plastid RPFs behave similarly to that of nucleus RPFs (**Figure 3A**, left). For 28-nt RPFs, their P-site is positioned 12 nt downstream from the 5′ end of the RPF, starting from the 13^th^ nt. As the RPF lengths decrease, the RPFs are shortened at the 5′ end, while the distance from the P-site to the 3′ end remains constant at 16 nt (including the first nt of the P-site) (**Figure 3A**, left). In contrast, the mitochondria P-site is located 7 nt downstream from the 5′ end for 28-nt RPFs, and the distance from the P-site to the 3′ end remains constant at 21 nt (**Figure 3A**, right).

**Figure 3.**
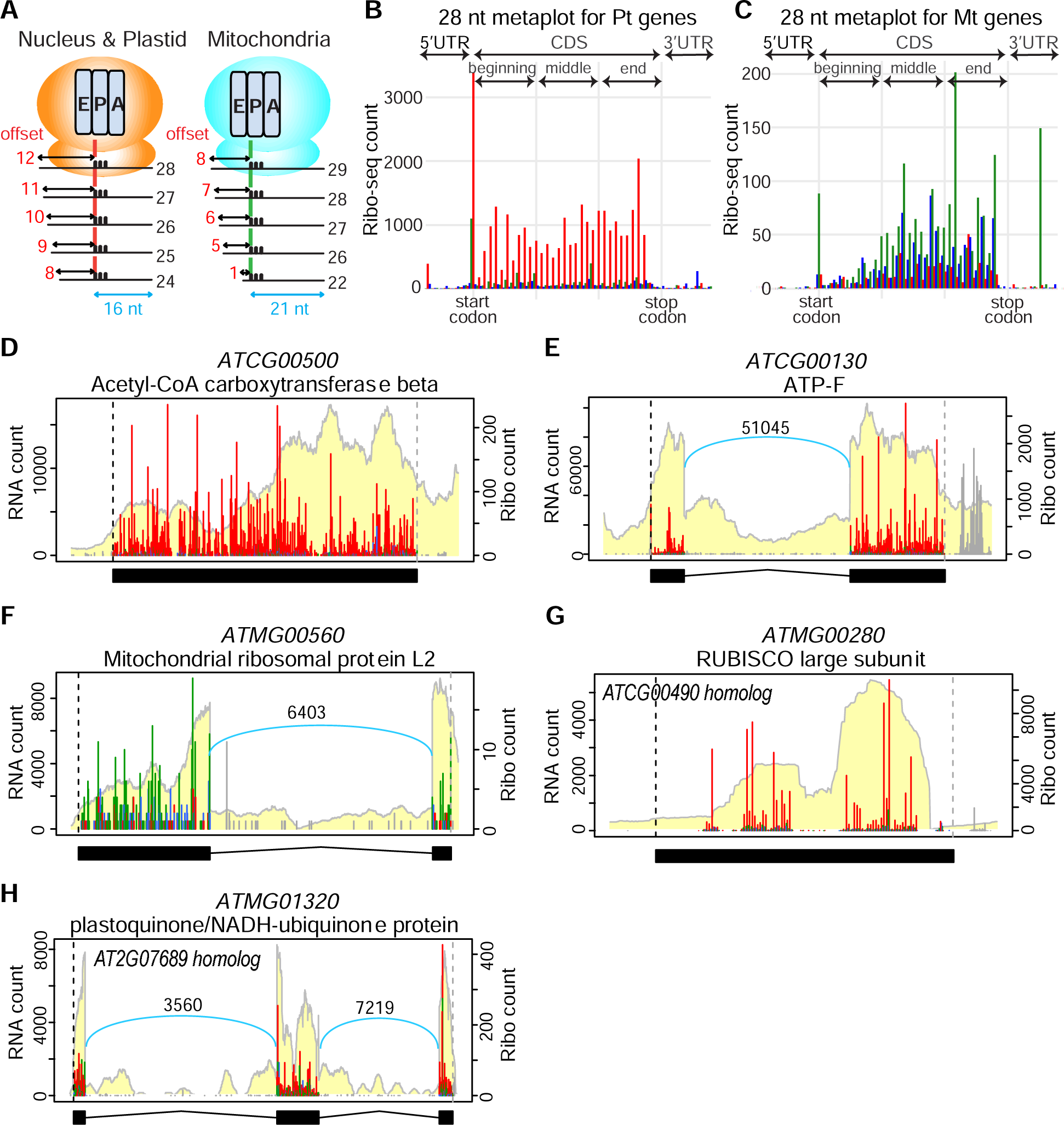
Translation in plastids and mitochondria. (A) P-site inferred in RPFs mapped to nucleus-, plastid- and mitochondria-encoded genes. (B–C) Metagene analysis of 28-nt RPFs at the start and stop codons of annotated plastid- and mitochondria-encoded ORFs. The RPFs are colored in red, blue, and green to indicate they are in the first, second, and third reading frames, respectively. (D–E) Examples of translational profiles of plastid-encoded genes, which use frame 1 (red). Note *ATCG00130* (E) contains an intron. (F) An example of translational profiles of a mitochondria-encoded gene, which uses frame 3 (green). Note it also contains an intron. (G–H) Two annotated mitochondria genes predicted to function in chloroplasts and have the RPFs mapped to frame 1 (red), suggesting these RPFs are from their plastid- and nucleus-encoded homologs, respectively, through multi-mapping. In (E, F, H), the number above the blue curly line indicates the read count across exon-exon junctions.

Consistently, metaplots show plastid RPFs exhibit similar patterns compared to nucleus RPFs, with strong 3-nt periodicity and enrichment of frame 1 (red) (**Figure 1A** and **3B**). However, plastid RPFs lack the signature blue signal (from frame 2) observed in nucleus RPFs at the codon preceding the stop (**Figure 1A and 3B**). In contrast, mitochondria RPFs show relatively weak 3-nt periodicity and mainly use frame 3 (green) (**Figure 3C**). Inspecting individual plastid and mitochondria genes also confirmed that plastid genes preferentially use frame 1 (red) (**Figure 3D–E**), while mitochondria genes predominantly use frame 3 (green) (**Figure 3F**). Interestingly, two mitochondria genes (*ATMG00280* and *ATMG01320*) were found to use frame 1 (red) (**Figure 3G–H**). Both genes were predicted to encode proteins that function in the chloroplasts: *ATMG00280* was predicted to encode RUBISCO large subunit, while *ATMG01320* was predicted to encode plastoquinone/NADH-ubiquinone protein. Coincidently, *ATMG00280* has a chloroplast-encoded homolog *ATCG00490*, and *ATMG01320* has a nucleus-encoded homolog *AT2G07689*. The usage of frame 1 (red) in these two mitochondria genes suggests that the RPFs detected were likely derived from their nucleus- and plastid-encoded homologs, respectively, and that *ATMG00280* and *ATMG01320* are not translated.

We observed substantial intron retentions in many plastid and mitochondria genes; however, the RPFs mainly mapped to the CDS/exon regions and are sparse in the intron regions (e.g., **Figure 3E–F**). These observations suggest that the translation occurs after the introns are spliced and that not all genes in plastids and mitochondria necessarily use a coupled transcription-translation mechanism like their prokaryotic ancestors (Xiong et al., 2022; Trösch, 2022).

Together these results demonstrate that our data are useful to validate the annotated gene models in plastids and mitochondria and to study the translation in these organelles.

### sORFs detected in tasiRNAs

Next, we investigated unconventional translation events on annotated ncRNAs, focusing on tasiRNAs. The biogenesis of tasiRNAs involves miRNA-guided cleavage of the primary transcripts of *TAS* transcripts (reviewed in (Fei et al., 2013)). There are four *TAS* gene families in Arabidopsis. Previous studies have reported the detection of RPFs on primary transcripts of *TAS1*, *TAS2*, and *TAS3*, especially at a sORF upstream of the miRNA target site (Li et al., 2016; Hsu et al., 2016; Bazin et al., 2017; Iwakawa et al., 2021). The RPF stalling associated with the sORF was found to be important for tasiRNA production (Iwakawa et al., 2021). However, it remains a question whether ribosomes actually translate these sORFs. Only the sORFs in *TAS3A* and *TAS4* have been detected significant 3-nt periodicity (Hsu et al., 2016; Hsu and Benfey, 2018). With the improved coverage in our dataset, RiboTaper identified actively translated ORFs across all four *TAS* gene families (**Figure 4 and 5**). It is noteworthy that translation occurs not only in the ORFs immediately adjacent to the miRNA target sites but also in ORFs upstream or downstream of the miRNA target sites.

**Figure 4.**
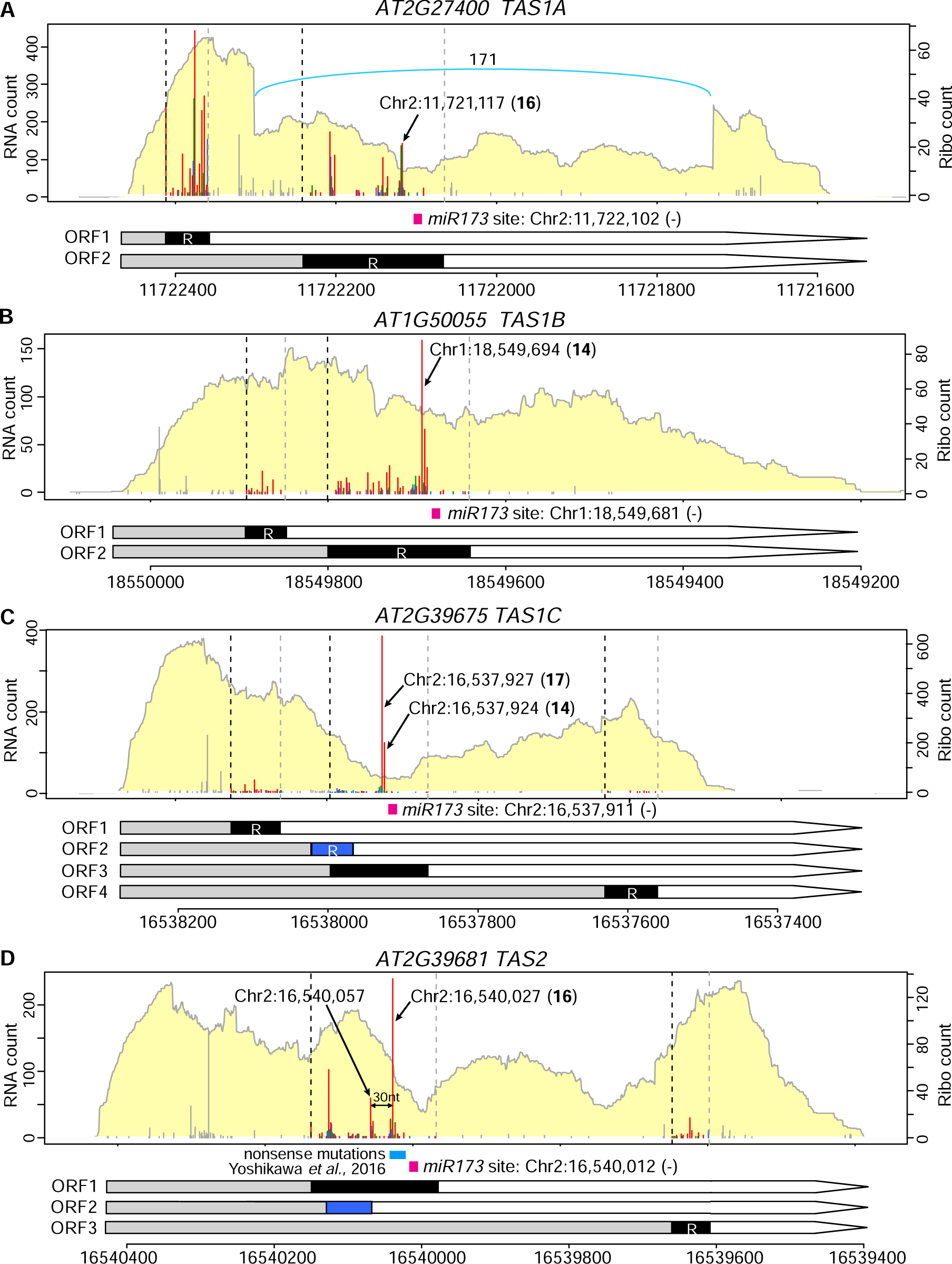
The translation and ribosome stalling of primary transcripts of *TAS1* and *TAS2.* (A–D) Expression profiles of *TAS1A–C* and *TAS2*. In the gene models, ORFs identified by RiboTaper are marked with ‘R.’ Above the gene models, miRNA target sites are indicated by hot pink rectangles. Note strong RPF peaks were observed upstream of the miRNA target sites (A-D). The coordinates and distance between the strong RPF peaks and the miRNA target sites are presented. The ORF3 in (C), and the ORF1 in (D) were manually curated based on their strong in-frame RPF peaks. These two ORFs were likely excluded by RiboTaper due to another ORF overlapping with them using a different reading frame (blue). Note the ORF1 in (D) was experimentally validated by a previous study (Yoshikawa et al., 2016). *TAS1A* contains an unannotated intron, which is indicated by a blue curly line, and the number above the blue curly line indicates the read count across exon-exon junctions.

**Figure 5.**
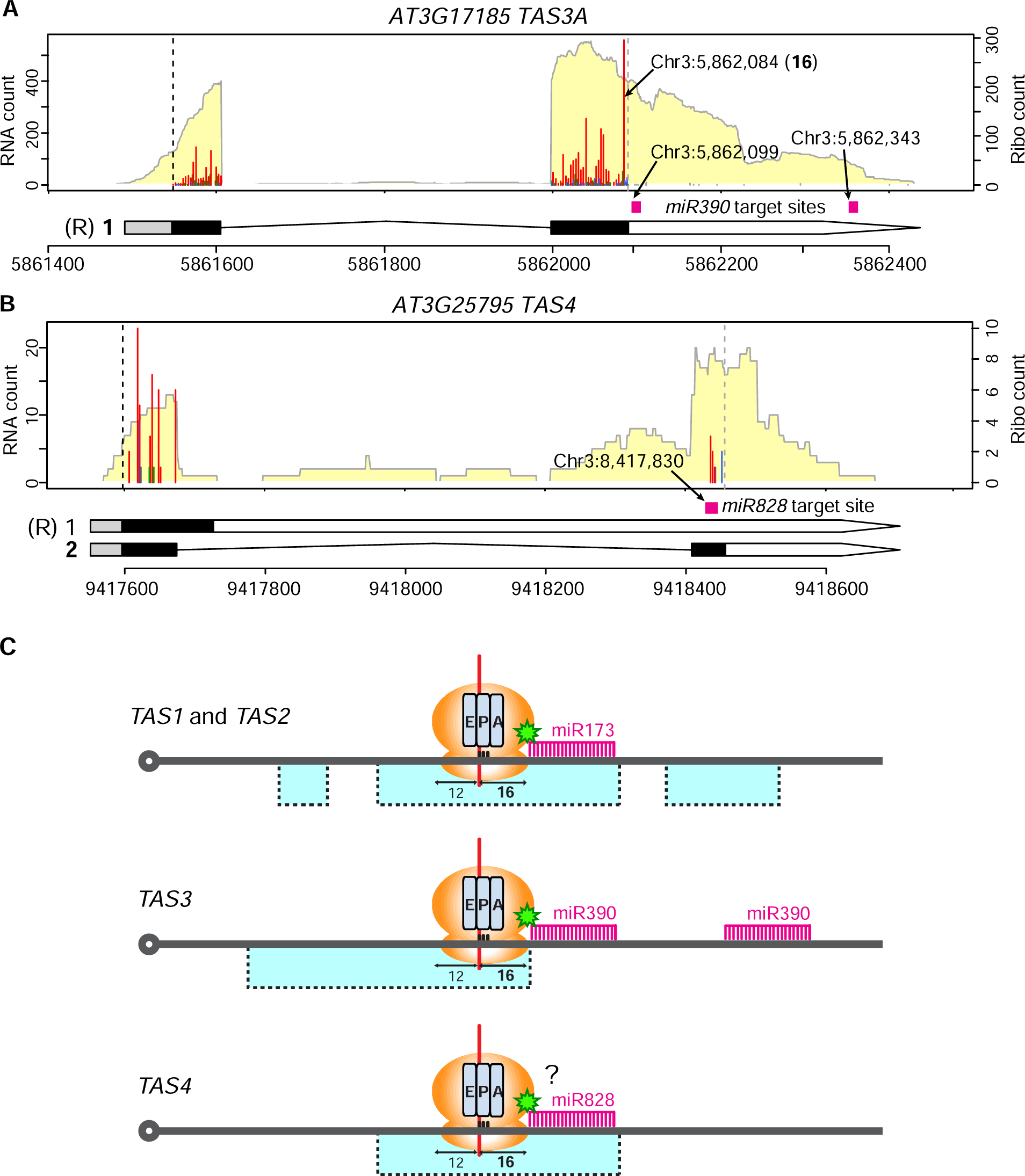
The translation of primary transcripts of *TAS3* and *TAS4* and the proposed models. (A–B) Expression profiles of *TAS3A* and *TAS4.* ORFs identified by RiboTaper were labeled with ‘(R)’ next to the gene models. Above the gene models, miRNA target sites are indicated by hot pink rectangles. In (A), a strong RPF peak was observed upstream of the first miRNA390 target site in *TAS3A*. The coordinates of the strong RPF peaks and the miRNA target sites are indicated. In (B), although RiboTaper identified an ORF within the only annotated transcript, our data revealed that *TAS4* has an additional isoform (isoform 2), and the ORF within this unannotated isoform is more likely translated. (C) Illustration of translated ORFs and RPF stalling relative to the miRNA target sites in *TAS1/2*, *TAS3*, and *TAS4*. In *TAS1/2*, multiple ORFs are translated. One particular ORF overlaps with the miR173 target site; the translation continues until the ribosome encounters the miR173 target site, where the first nt within the ribosomal P-site (red vertical line) corresponds to 16 nt upstream of the miR173 target site. In *TAS3*, only one ORF is translated, and the ORF is entirely upstream of the first miR390 target site. Similarly, the translation continues until the ribosome encounters the first miR390 target site, where the ribosomal P-site corresponds to 16 nt upstream of the miR390 target site. In *TAS4*, only one ORF is translated, and the ORF overlaps with the miR828 target site. Light blue boxes indicate ORFs.

In *TAS1A*, two ORFs were identified by RiboTaper (**Figure 4A**). Interestingly, *TAS1A* contains an unannotated intron, suggesting that *TAS1A* could also be regulated by alternative splicing. ORF2 is located within this retained intron and overlaps with the miR173 target site (**Figure 4A**). Importantly, RPF stalling was observed 16 nt upstream of the miR173 target site (**Figure 4A**). Similarly, in *TAS1B*, two ORFs were detected by RiboTaper, and strong RPF stalling was observed 14 nt upstream of the miR173 target site (**Figure 4B**). *TAS1C* has three ORFs identified by RiboTaper (ORF1, 2, and 4). Additionally, another ORF (ORF3) overlaps with the miR173 target site, and strong RPF pausing signals were detected within this ORF at 14 and 17 nt upstream of the miR173 target site (**Figure 4C**). It should be noted that ORF3 was likely excluded by RiboTaper due to the overlapping ORF2, which utilizes a blue reading frame (**Figure 4C**).

In *TAS2*, strong 3-nt periodicity was observed in the ORF1, accompanied by distinct RPF stalling 16 nt upstream of the miR173 target site (**Figure 4D**). Another strong pausing was observed 30 nt upstream of the pausing adjacent to the miR173 target site.

Unlike *TAS1* and *TAS2*, *TAS3A* possesses two miR390 target sites. In line with our previous findings (Hsu et al., 2016; Hsu and Benfey, 2018), the ORF located entirely upstream of the first miR390 target site was identified by RiboTaper and exhibits strong 3-nt periodicity (**Figure 5A**). Moreover, strong stalling was detected 16 nt upstream of the first miR390 target site (**Figure 5A**).

*TAS4* has relatively low expression in our data. Nevertheless, consistent with our previous findings (Hsu et al., 2016), RiboTaper detected an ORF in the only annotated transcript (isoform 1). However, when we visualized the RNA-seq and Ribo-seq profiles, we realized that *TAS4* has an unannotated isoform (isoform 2). The RPFs located within the ORF in isoform 2 overlap with the miR828 target site (**Figure 5B**).

Together our data provide evidence of extensive RPFs associated with *TAS1*, *TAS2*, *TAS3*, and *TAS4*. The significant 3-nt periodicity observed in these ORFs strongly supports their active translation. In a previous study on *TAS2*, it was shown that the position of ORF termination relative to the miR173 target site is crucial for tasiRNA biogenesis (Yoshikawa et al., 2016). In *TAS1*, *TAS2*, and *TAS3*, we observed that RPFs accumulate until 14-17 nt upstream of the miRNA target site. Considering that our RPFs were plotted using the first nt of the P-site (the 13^th^ nt for 28-nt RPFs) and the distance between the P-site to the 3′ end remains constant at 16 nt (**Figure 3A**), the observed distance between the RPF pausing and the miRNA target site coincides with the approximate length of the RPF from the P-site to the 3’ end (illustrations in **Figure 5C**). In line with the previously proposed ‘ribosome stalling complex’ model (Iwakawa et al., 2021), our observations suggest that most ribosomes can translate until they encounter the miRNA/AGO/SGS3 complex, leading to ribosome stalling, in *TAS1*, *TAS2*, and *TAS3* (**Figure 5C**). These results are also consistent with the previous finding that a mutation within 6 nts upstream of the miR173 target site does not have observable effects (Yoshikawa et al., 2016), as the last ribosomal P-site is 16 nt upstream of the miRNA target site (**Figure 5C**). Thus, our improved coverage data enhance the resolution and identification of translated ORFs on *TAS* transcripts.

### Translation of dORFs

Next, we examined unannotated ORFs on protein-coding genes. In our dataset, RiboTaper identified 209 dORFs (**Figure 2A** and **2B**). This class of ORFs has not been described in detail in plants, and their functional significance remains unknown. Interestingly, we found that the mORFs of genes containing dORFs exhibit higher translation efficiency compared to genes without dORFs, suggesting that dORFs are associated with high translation efficiency genes (**Figure 6A**). Indeed, the gene ontology (GO) term of the dORF-containing genes is enriched for genes expected to be highly expressed and translated, such as ribosome and translation machinery (**Dataset S4**).

**Figure 6.**
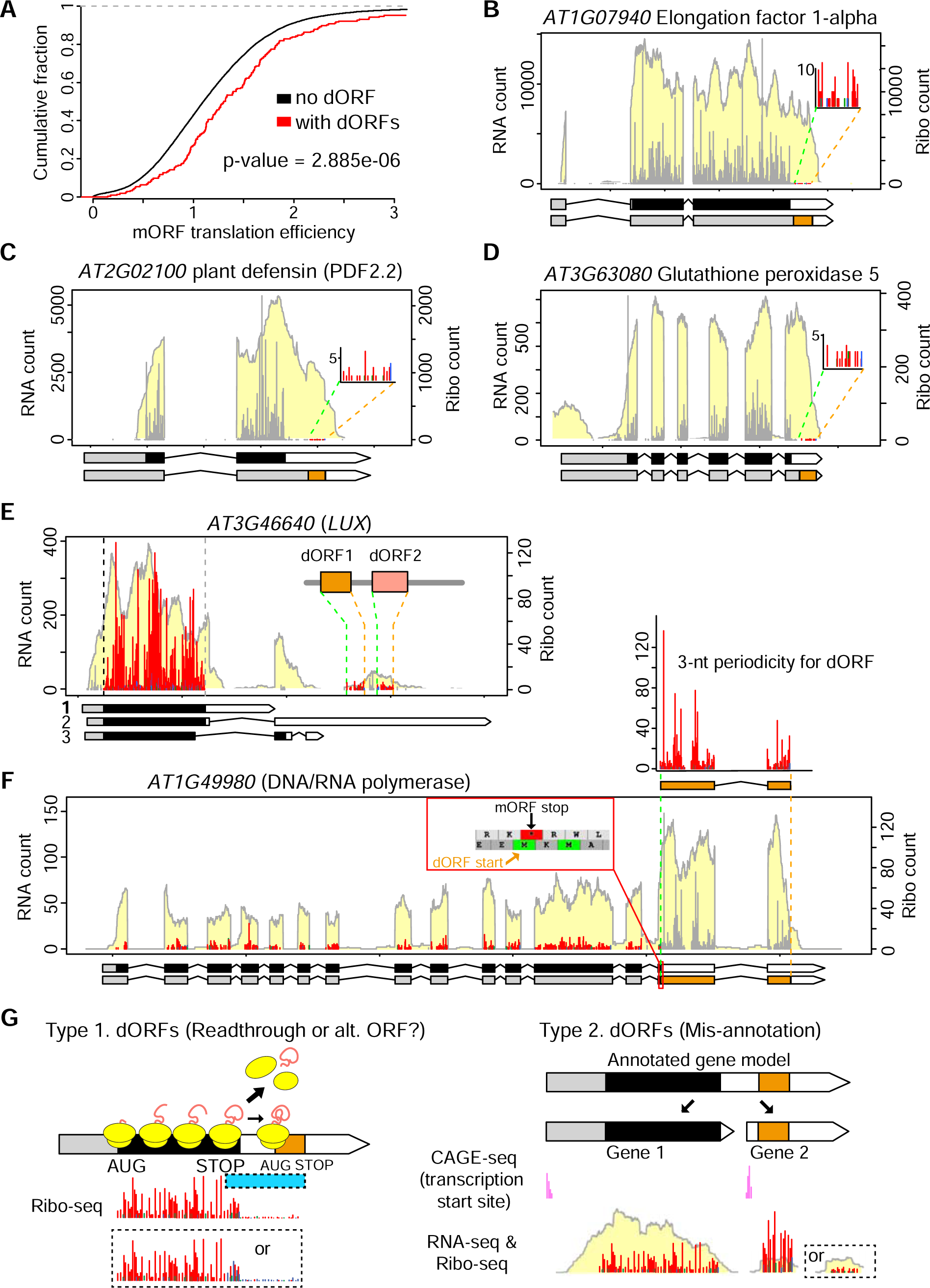
Examples of dORFs and their translation efficiency. (A) Comparison of mORF translation efficiency between genes with and without dORFs. The statistical significance was determined by the Kolmogorov–Smirnov test. (B–F) Examples of dORFs. In the gene models, the dORFs are indicated by orange box(es), and their translation start and stop are indicated by green and orange vertical dashed lines, respectively, within the profiles. (B–D) Type 1 dORFs: the mORF have high translation levels, while the dORFs have low translation levels. The RPFs in the mORFs are shown in grey. The RPFs in the dORFs are magnified and presented in three colors to indicate their reading frame. (E–F) Type 2 dORFs: an additional gene/transcript is present in the annotated 3′ UTR or downstream of the mORF gene. These additional genes/transcripts have distinct mRNA levels compared to the mORF genes. The zoom-in in (F) shows the dORF start overlaps with the mORF stop, supporting these two ORFs are unlikely to be translated sequentially on the same mRNA. (G) Illustrations of Type 1 and Type 2 dORFs. Type 1 dORFs could be potential readthrough from the mORF, or continuous translation from an alternative ORF overlapping with the mORF. Type 2 dORFs result from mis-annotation, in which a hidden gene/transcript is located in the 3′ UTR or downstream of the mORF gene. These hidden genes have independent transcription start sites supported by published CAGE data (see **Figure S7**), and they have distinct RNA-seq (and Ribo-seq) levels, compared to the upstream mORF.

We further evaluated these dORFs by analyzing their translational profiles (**Figure 6B– F**). While we confirmed these dORFs display strong 3-nt periodicity, we observed that most of the dORFs have low translation levels (**Figure 6B–E**). Notably, their patterns could be categorized into two groups, suggesting that they are unlikely to be reinitiation events following the termination of the mORFs. In Type 1 dORFs, the mORF has relatively high translation levels (**Figure 6B–D**), and substantial RPFs are present between the mORF and the dORF (**Figure S6A-C**). This implies that the RPFs in the dORF could result from a readthrough of the mORF using frame 1 (red) (**Figure S6A**) or continued translation from an alternative ORF using a different frame (e.g., blue, **Figure S6B–C**).

The Type 2 dORFs appear to originate from unannotated small genes/transcripts that are located downstream or overlapping with the 3′ UTRs of the annotated genes. These dORFs are likely derived from independent genes/transcripts, as their mRNA levels differ from those of the annotated genes (**Figure 6E–F**). For example, there is a small gene that was annotated as part of the 3′ UTR of isoform 2 of *AT3G46640 LUX* (a key transcription factor in the circadian clock), and RiboTaper identified two translated ORFs encoding 58 and 54 aa in this unannotated gene (**Figure 6E**). Similarly, a dORF was identified in the 3′ UTR of *AT1G49980* (**Figure 6F**). The start codon of the dORF overlaps with the stop codon of the mORF (zoomed-in section in **Figure 6F**), indicating these two ORFs cannot be translated sequentially on the same mRNA. Published CAGE data (Thieffry et al., 2020) revealed additional transcription start sites associated with these dORFs (**Figure S7A–B**), providing further support that these unannotated ORFs, hidden within the 3′ UTRs or downstream regions of other genes, originate from independent genes/transcripts.

Interestingly, a tBLASTn search of the 153 dORFs greater than 20 aa revealed that 11 of them are conserved throughout evolution (**Figure S8**). Importantly, all 11 conserved dORFs belong to the Type 2 dORFs (**Figure S8**). This finding suggests that the peptide sequences encoded by these unannotated genes/dORFs likely correspond to functional proteins.

Taken together, the dORFs we detected are unlikely to be reinitiation events analogous to the uORF-mORF pairs. The Type 1 dORFs could potentially result from readthrough events or part of alternative ORFs (**Figure 6G, left**); further investigation is needed to elucidate the mechanisms. On the other hand, the Type 2 dORFs revealed the presence of unannotated small genes hidden in the 3′ region of other genes (**Figure 6G, right**).

### Ribosome queueing at CPuORFs links to co-translational decay

We next investigated uORFs, which represent the largest class of unannotated ORFs we identified (**Figure 2A**). CPuORFs are of great interest as they are associated with significant ribosome pausing/repression, likely through the interaction between the nascent peptide and the exit tunnel within ribosomes. Notably, several CPuORFs have been reported to interact with certain metabolites (reviewed in (Van Der Horst et al., 2020)). CPuORFs are also a focus of co-translational decay studies (Yu et al., 2016; Hou et al., 2016). Co-translational decay demonstrates that mRNAs can undergo degradation while bound by translating ribosomes. As a result, the degraded mRNA fragments display features associated with translation (Hu et al., 2009; Pelechano et al., 2015; Merret et al., 2015; Yu et al., 2016; Hou et al., 2016). Global analysis of 5′ ends of mRNA degradation fragments in Arabidopsis detected mRNA fragments with ∼30-nt periodicity (approximate length of RPFs) preceding the stop codons of several CPuORFs, supporting the idea that several ribosomes tightly stack behind a pausing ribosome in these regions (Yu et al., 2016; Hou et al., 2016). However, the corresponding 30-nt periodicity of RPFs was not detected in previous Ribo-seq data (Hou et al., 2016).

In Arabidopsis, 119 CPuORFs have been identified and they are grouped into 81 homology groups based on the conservation of the uORF peptide sequences (reviewed in (Van Der Horst et al., 2020)). We examined the RPFs in different homology groups of CPuORFs and found those in Homology Group 1 showed clear periodic ribosome pausing patterns. Specifically, 2–4 distinct RPF peak(s) at 30-nt intervals were observed upstream of the stop codons of CPuORF1, 2 and 3, within *bZIP2*, *bZIP11*, and *bZIP53* transcripts, respectively (**Figure 7A–C**). These patterns are consistent with the expectation that multiple ribosomes line up behind a pausing ribosome prior to the stop codon of these CPuORFs (see illustrations of ribosome positions above the RPF peaks in **Figure 7A–C**). We also observed a similar pattern in *MYB51* (**Figure 7D**), which contains a Brassicaceae-specific CPuORF (Hou et al., 2016). Therefore, our enhanced Ribo-seq data provide evidence for the periodic ribosome pausing upstream of the stop codons of the CPuORFs, further supporting the occurrence of co-translational decay in plants.

**Figure 7.**
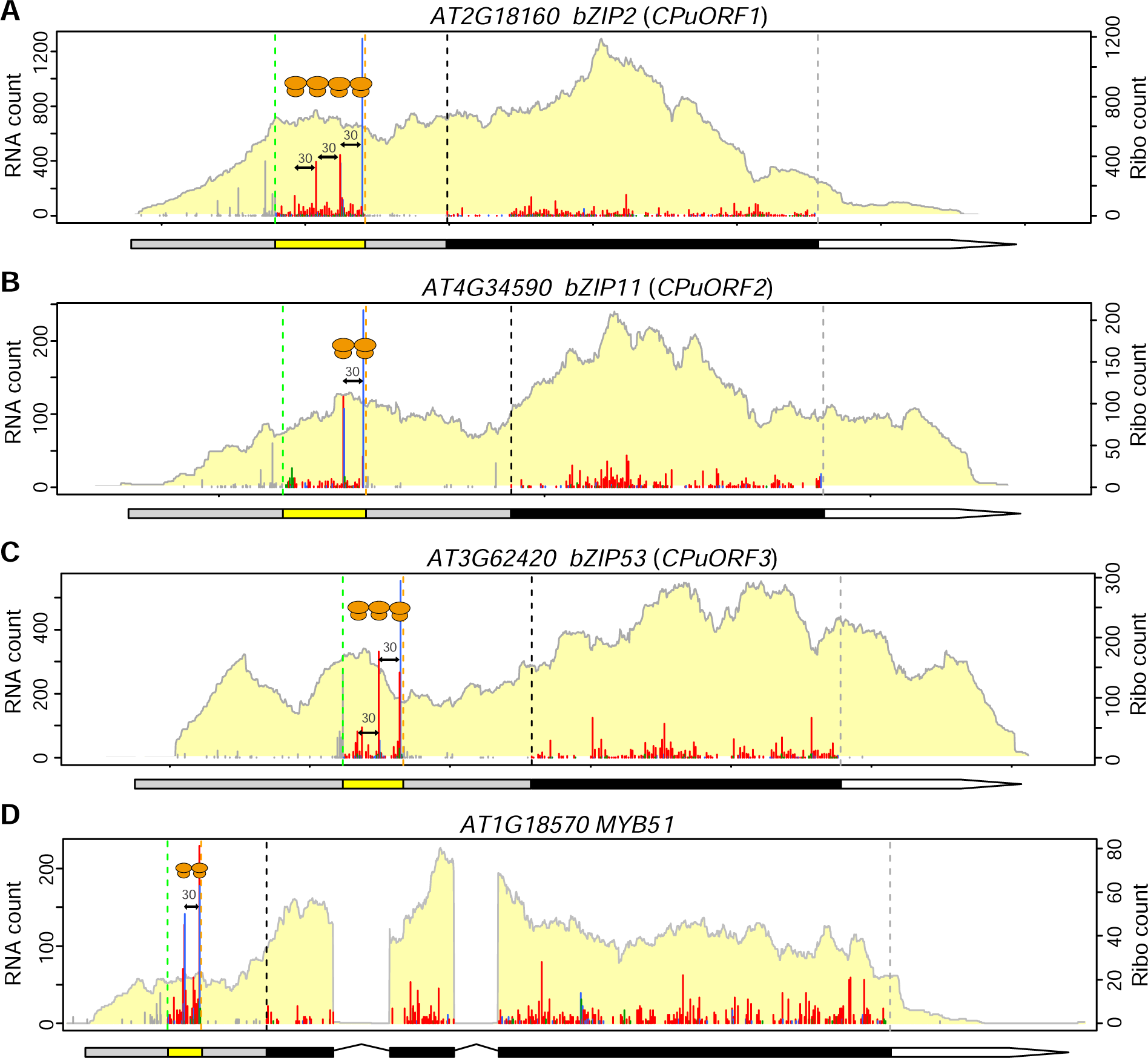
Periodic ribosome stalling in CPuORFs. (A-D) Examples of CPuORFs with periodic ribosome stalling. In the gene model, CPuORF is shown by a yellow box, and their translation start and stop are indicated by green and orange vertical dashed lines, respectively, within the profiles. The 30-nt internals between RPF peaks are indicated, and the inferred ribosome positions are illustrated.

### Identification of uORFs using RiboTaper

Besides CPuORFs identified by evolutionary conservation, we also investigated uORFs identified through our experimental approach. uORFs represent the shortest unannotated ORFs (**Figure 2E**), and their identification could be most impacted by the RPF coverage. Our new data significantly improved the detection of uORFs based on the 3-nt periodicity, increasing the number from the previous 187 uORFs to 2113 uORFs (**Figure 2A–B**, **Dataset S1A**). Strong 3-nt periodicity within the uORF regions in the translational profiles supports the active translation of these uORFs (examples shown for *S6K1* and *RGA1* in **Figure 8A–B**). Comparing uORF genes with similar mRNA levels in our previous and current datasets, it is evident that both the uORF and mORF regions exhibit improved coverage in our current data (**Figure 8A–B**).

**Figure 8.**
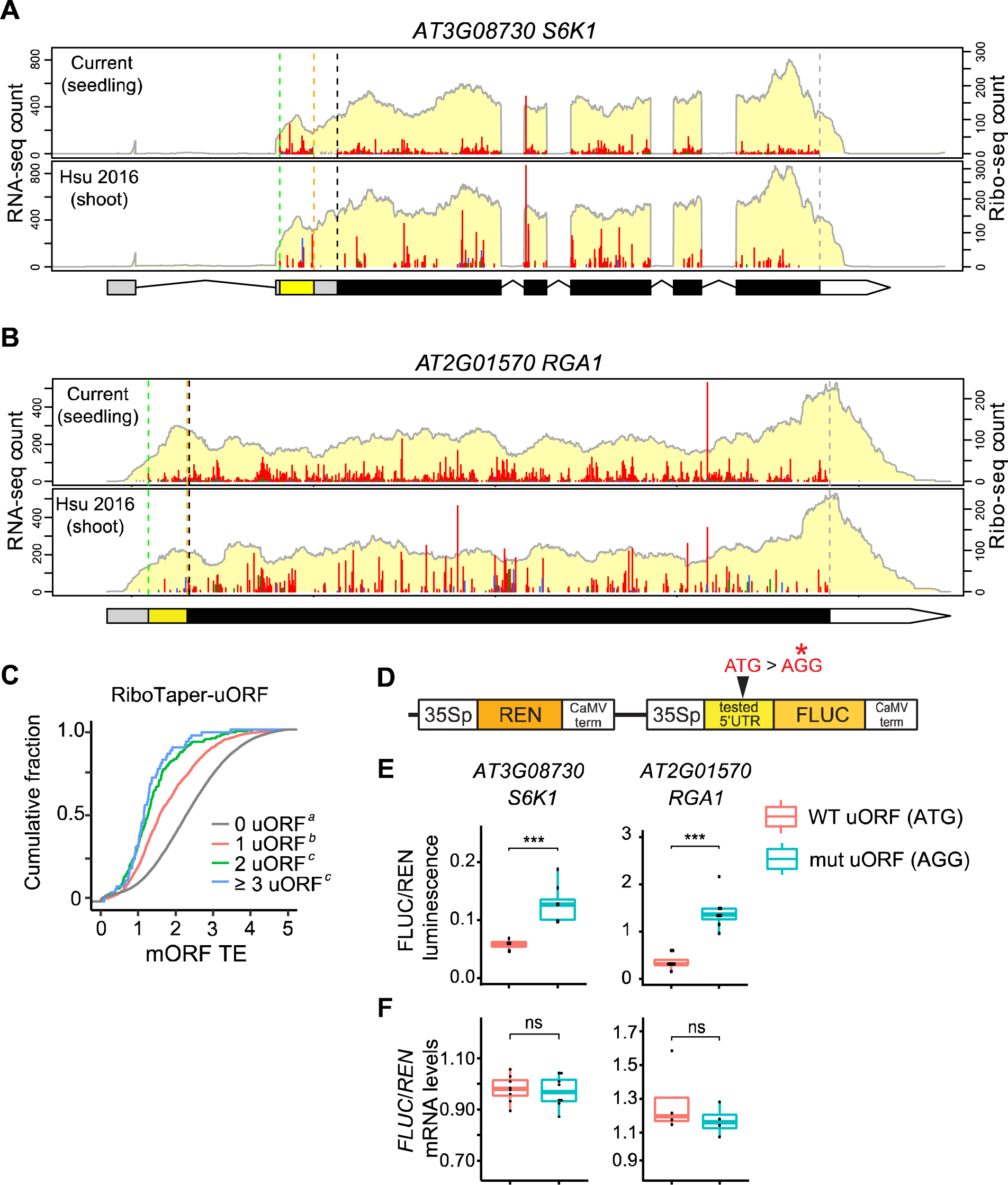
Newly discovered uORFs by RiboTaper. (A–B) Two examples of translated uORFs identified by RiboTaper in the current data compared to our previous data (Hsu et al., 2016). (C) Cumulative plot comparing the mORF translation efficiency (TE) of genes containing 0, 1, 2, ≥ 3 translated uORFs identified by RiboTaper. The different italic superscript letters indicate the statistical significance between groups (Kolmogorov–Smirnov test, p < 0.05). (D) The design of dual-luciferase constructs for testing uORF functions. The start codon of uORF (ATG) is mutated to AGG in the mutated version. 35Sp: 35S promoter; CaMV term: CaMV terminator. (E) Relative FLUC luminescence comparing 5′ UTRs carrying the wild-type uORF or mutated uORF. FLUC luminescence levels are normalized to REN luminescence levels. (E) Relative *FLUC* mRNA levels comparing 5′ UTRs carrying the wild-type uORF or mutated uORF. *FLUC* mRNA levels are normalized to *REN* mRNA levels. The statistical significance for boxplots in (E–F) was determined by Wilcoxon rank sum test (*: 0.01 < p < 0.05, **: 0.001 < p < 0.01, ***: 1e-4 < p < 0.001).

Consistent with our expectations, these translated uORFs are associated with lower mORF translation efficiency (**Figure 8C**). Furthermore, genes that have a higher number of translated uORFs exhibit lower mORF translation efficiency (**Figure 8C**).

To validate that these uORFs function as translational repressors, we performed a dual-luciferase assay comparing wild-type uORFs with mutated uORFs, where the start codon ATG was mutated to AGG, in *S6K1* and *RGA1* (**Figure 8D** and **8E**). Upon mutating the uORFs, the downstream FLUC reporter showed increased protein levels/activities (**Figure 8E**), while the *FLUC* transcript levels showed no significant changes (**Figure 8F**). These results suggest that these uORFs normally repress the translation of the downstream mORFs.

### Identification of uORFs using CiPS

Although the uORFs identified by RiboTaper show significant 3-nt periodicity, we observed that globally, uORFs have the lowest 3-nt periodicity compared to CCDSs and ncORFs (**Figure 9A**), and the 3-nt periodicity appears to be inversely correlated with the ORF length among these ORF groups (**Figure 9A** and **Figure 2E**). We reasoned that the overall 3-nt periodicity is affected by the atypical RPFs observed at the codon proceeding the stop codon (refer to as ‘(−1) codon’ hereafter). In this position, the P-sites of many RPFs are mapped to frame 2 (blue) instead of the expected frame (red) (**Figure 1A** and illustrations in **Figure 9B–E**). This pattern likely arises from the binding of release factors, rather than a charged tRNA, to the A-site during termination, leading to a different ribosomal conformation and resulting in distinct RPF patterns (Alkalaeva et al., 2006; Brown et al., 2015; Matheisl et al., 2015). In addition, given termination is a major rate-limiting step during translation, more RPFs are associated with this position (Wolin and Walter, 1988; Ingolia et al., 2009). Importantly, this out-of-frame signal (blue) can disproportionally and significantly reduce the 3-nt periodicity of short ORFs and impact their identification based on 3-nt periodicity (illustrations in **Figure 9B–E**). Consistently, comparing the size distribution of predicted uORFs (i.e., AUG-initiated ORFs in the 5’ UTR) and those identified by RiboTaper reveals that RiboTaper clearly biases towards longer uORFs (**Figure 9F**, median 9 vs. 21, respectively).

**Figure 9.**
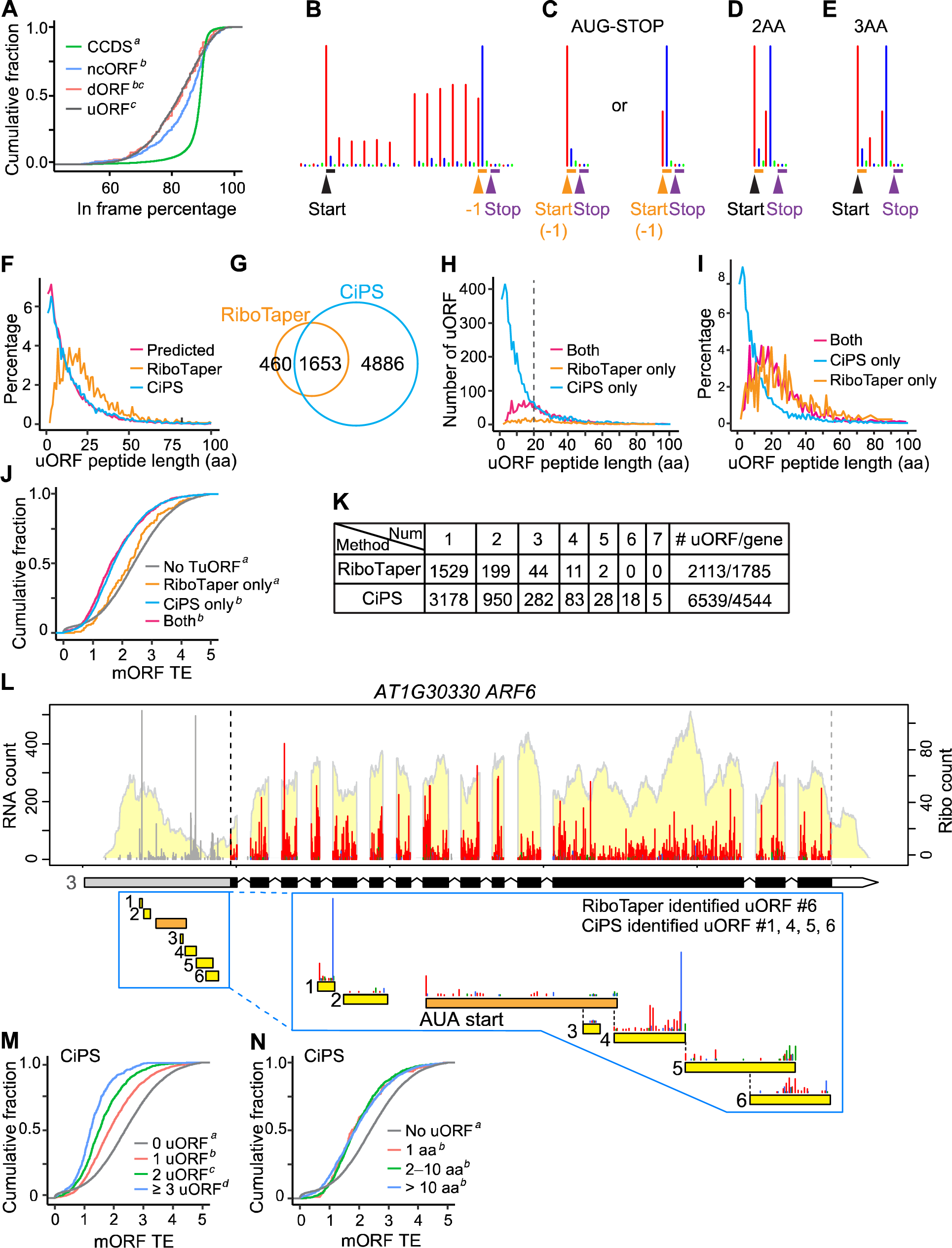
Identification of uORFs using CiPS. (A) Cumulative plot comparing the in-frame percentage of CCDSs, ncORFs, dORFs, and uORFs identified by RiboTaper. The different italic superscript letters indicate the statistical significance between groups (Kolmogorov–Smirnov test, p < 0.05). (B–E) Illustrations of expected RPF distribution of a long uORF (B), a minimum uORF (C), a 2-aa uORF (D), and a 3-aa uORF (E). Note that the codon preceding the stop (−1) is expected to have a significant amount of RPFs mapped to frame 2 (blue). For minimum uORFs (C), the start codon is also the (−1) codon. (F) Distribution of uORF peptide length of predicted uORFs (AUG-start), RiboTaper-identified uORFs, and CiPS-identified uORFs. (G) Venn diagram for CiPS and RiboTaper identified uORFs. (H) Distribution of uORF peptide length, in terms of number, comparing uORFs identified by RiboTaper-only, CiPS-only, or both methods. (I) Distribution of uORF peptide length, in terms of percentage, comparing uORFs identified by RiboTaper-only, CiPS-only, or both methods. (J) Cumulative plot comparing mORF translation efficiency of genes without any translated uORFs and with uORFs identified by RiboTaper-only, CiPS-only, or both methods. The different italic superscript letters indicate the statistical significance between groups (Kolmogorov– Smirnov test, p < 0.05). (K) Comparison of the number of translated uORFs identified per gene between RiboTaper and CiPS. (L) The expression profile of *ARF6*, which contains multiple uORFs. Zoom-ins of each AUG-uORF (yellow box) and an AUA-start uORF (orange box) are presented to evaluate the performance of RiboTaper and CiPS. (M) Cumulative plot comparing the mORF translation efficiency of genes containing 0, 1, 2, ≥ 3 translated uORFs identified by CiPS. The different italic superscript letters indicate the statistical significance between groups (Kolmogorov–Smirnov test, p < 0.05). (N) Cumulative plot comparing the mORF translation efficiency of genes containing no, 1 aa (minimum), 2–10 aa, and > 10 aa translated uORFs identified by CiPS. The different italic superscript letters indicate the statistical significance between groups (Kolmogorov–Smirnov test, p < 0.05).

To improve the identification of short uORFs, we developed a new approach called CiPS (<u>C</u>ount, in-frame <u>P</u>ercentage and <u>S</u>ite). In brief, we first classified the blue signal at the (−1) codon as in frame. We then applied three criteria to consider if a uORF is translated by evaluating (1) RPF counts, (2) in-frame RPF percentage, and (3) in-frame sites occupied by RPFs (see Materials and Methods for detail).

In total, CiPS identified 6539 translated uORFs (**Dataset S5A**), including 370 minimum uORFs (1 aa) and 388 2-aa uORFs. In contrast, RiboTaper identified no minimum uORFs and only one 2-aa uORF (**Dataset S1A**). CiPS successfully identified a significant number of short uORFs (**Figure 9H–I**). Notably, the uORFs identified by CiPS exhibit a similar size distribution compared to the predicted uORFs (**Figure 9F**). Comparing the uORFs identified by RiboTaper and CiPS, 1653 uORFs were identified by both methods, while 460 uORFs were only found by RiboTaper and 4886 were only found by CiPS (**Figure 9G**). The uORFs identified exclusively by RiboTaper have a marginal effect on the translation efficiency of their mORFs (**Figure 9J**). In contrast, the uORFs identified by CiPS only or by both methods show significantly stronger repression (**Figure 9J**).

CiPS also improved the number of uORFs identified within individual transcripts, with the highest number of 7 uORFs detected within a transcript (**Figure 9K**). We manually plotted a set of genes to compare the performance of RiboTaper and CiPS. In the example of *ARF6* (**Figure 9L**), there are six potential AUG-initiated uORFs. CiPS identified uORF-1, 4, 5, and 6. In contrast, RiboTaper only identified uORF-6. By chance, we also observed a noncanonical uORF initiated by an AUA start codon (**Figure 9L**). uORF-1 and 4 were excluded by RiboTaper, likely due to the strong blue signal at the (−1) codon, as well as the partial overlap between uORF-4 and neighboring uORFs (**Figure 9L**).

Consistent with our expectations and the findings with RiboTaper (**Figure 8C**), genes containing a higher number of CiPS-identified uORFs show a correlation with lower mORF translation efficiency (**Figure 9M**).

A previous survey of 28 plant uORFs in the literature suggested that longer uORFs are associated with stronger repression of mORFs (Von Arnim et al., 2014). We investigated whether there is a relationship between uORF length and repression with our genome-wide dataset. We compared genes with one translated uORF encoding 1, 2–10, or > 10 aa, and found they similarly repress the translation efficiency of the mORFs, compared to genes without any translated uORFs (**Figure 9N**). This result demonstrates these short uORFs are equally powerful translational repressors.

### Characterization of minimum uORFs and tiny uORFs

By definition, ‘AUG-stop’ represents the shortest possible uORF, known as the minimum uORF (illustrations in **Figure 9C**). In our dataset, 746 minimum uORFs contain at least one RPF. As minimum uORFs only have 1 codon, we modified the in-frame site criteria of CiPS accordingly (see Materials and Methods for details) and uncovered 370 minimum uORFs passed the criteria (**Dataset S5B**). This list includes the two minimum uORFs previously reported in *NIP5;1* (**Figure 10A**), and the minimum uORFs reported in *SKU5* and *BOR2* (**Figure 10B–C**), which are related to boron transport and response (Tanaka et al., 2016; Sotta et al., 2021). Our data did not detect the reported *ABS2*, as in our experimental conditions, *ABS2* uses a transcription start site downstream of the potential minimum uORF (**Figure S9**). Notably, we detected new minimum uORFs in several important genes, including *PBL36*, a receptor-like kinase involved in regulating shoot and root meristems; *SnRK2.2*, a key kinase in the ABA signaling pathway; and *NPR3*, a salicylic acid receptor and a negative regulator of plant pathogen response (**Figure 10D–F**). As expected, these minimum uORFs show the characteristic blue signal at the (−1) codon, since minimum uORFs consist of only one codon, which is also immediately upstream of the stop codon (**Figure 10A–F**, illustrations in **Figure 9C**).

**Figure 10.**
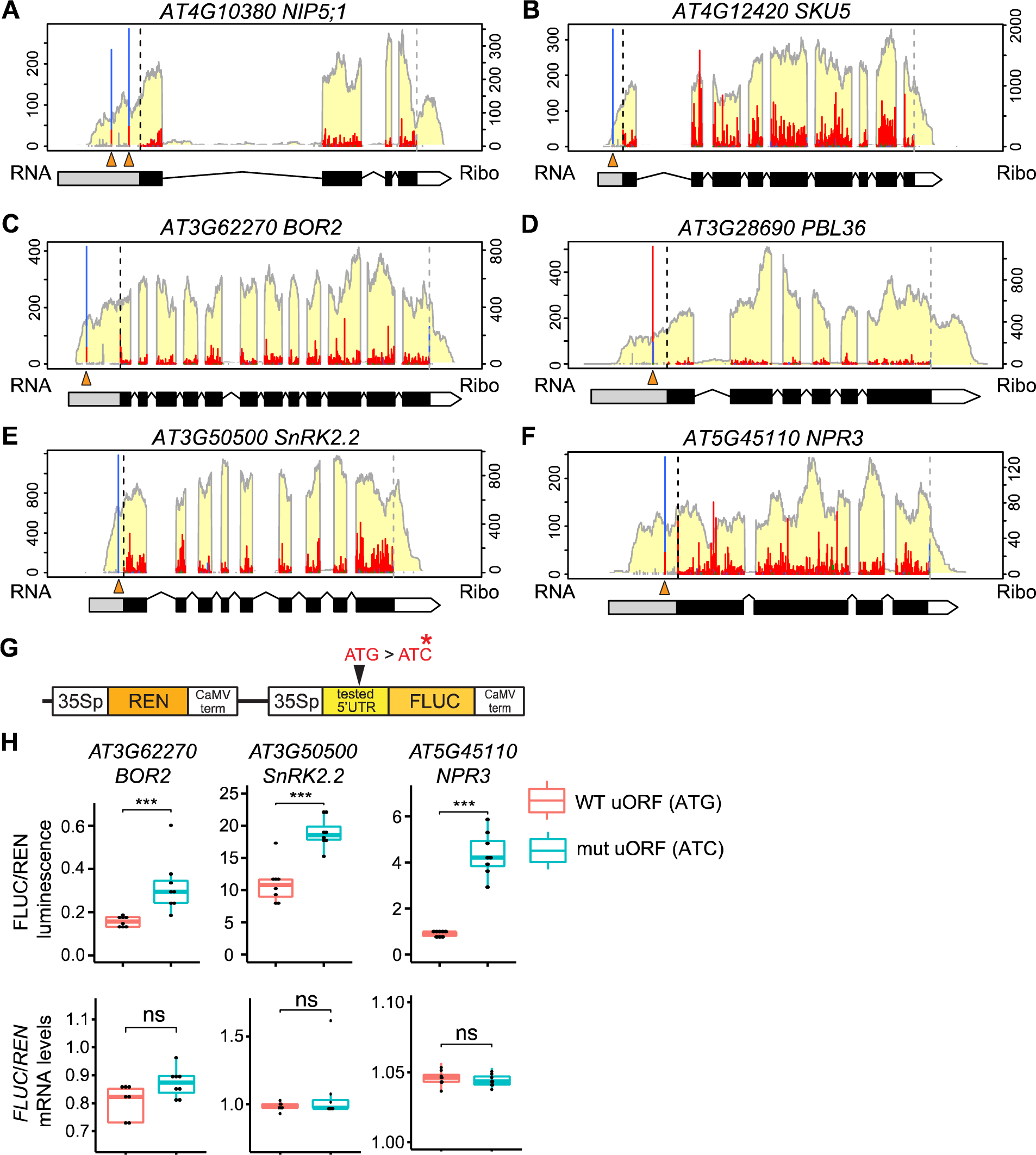
Translation of minimum uORFs. (A–F) Examples of minimum uORFs. The minimum uORF positions are indicated by orange triangles. RPFs mapped to minimum uORFs show the characteristic of (−1) codon, where a high fraction of RPFs mapped to frame 2 (blue). (G) The design of dual-luciferase construct testing minimum uORF functions. The start codon of minimum uORF (ATG) is mutated to ATC in the mutated version. (H) Relative FLUC luminescence comparing 5′ UTRs carrying the wild-type uORF or mutated uORF. FLUC luminescence levels are normalized to REN luminescence levels. (I) Relative *FLUC* mRNA levels comparing 5′ UTRs carrying the wild-type uORF or mutated uORF. *FLUC* mRNA levels are normalized to *REN* mRNA levels. The statistical significance for boxplots in (H–I) was determined by Wilcoxon rank sum test (*: 0.01 < p < 0.05, **: 0.001 < p < 0.01, ***: 1e-4 < p < 0.001).

As shown in **Figure 9N**, globally, minimum uORFs show similar repression effects on downstream mORF translation, compared to longer uORFs. To further validate the functions of these minimum uORFs, we performed the dual-luciferase assay to test the uORFs in *BOR2*, *SnRK2.2* and *NPR3* (**Figure 10G** and **10H**). We found that mutations in the uORFs lead to increased FLUC levels without significantly affecting *FLUC* transcript levels (**Figure 10H** and **10I**). These results support that these minimum uORFs normally repress their mORF translation.

Like minimum uORFs, other uORFs with extremely short lengths are difficult to be identified based on 3-nt periodicity (illustrations in **Figure 9D–E**). CiPS successfully identified 2984 tiny uORFs (2–10 aa) (**Dataset S5C**). Examples of 2-, 3-, 4-, 5-, and 8-aa uORFs are shown in **Figure 11A-D** and **S10A–B**. Similar to the minimum uORFs, mutations in these tiny uORFs result in increased FLUC levels without significantly increasing the *FLUC* transcript levels (**Figure 11E, S11** and **S10C–D**). These results support that these tiny uORFs normally repress their mORF translation and are consistent with their repressive effects observed in global analysis (**Figure 9N**).

**Figure 11.**
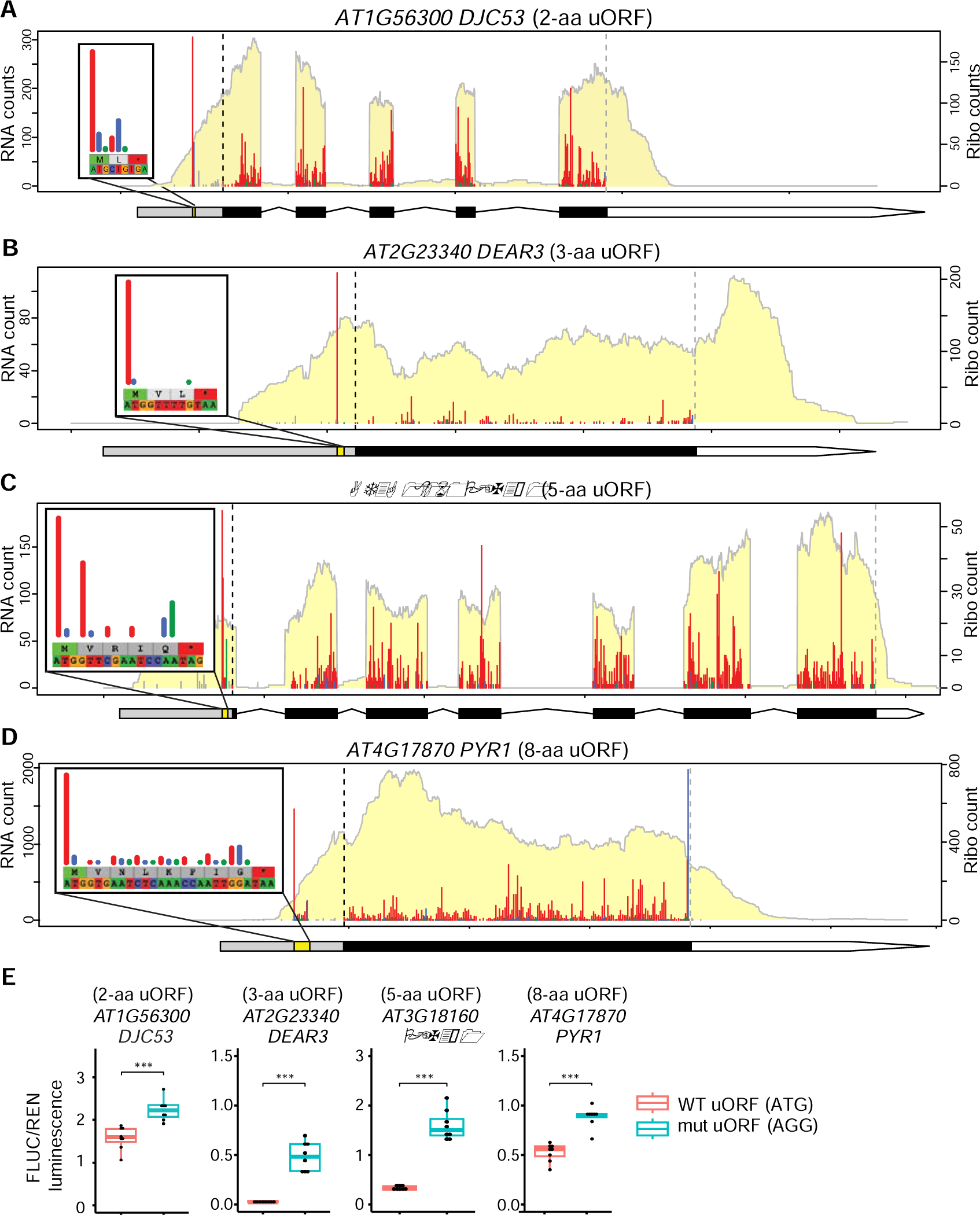
Translation of tiny uORFs. (A–D) Examples of tiny uORFs of varying uORF length. The Ribo-seq read and frame information for the tiny uORFs are shown in the boxes. (E) Relative FLUC luminescence comparing 5′ UTRs carrying the wild-type uORF (ATG) or mutated uORF (AGG). FLUC luminescence levels are normalized to REN luminescence levels. Statistical significance was determined by Wilcoxon rank sum test (*: 0.01 < p < 0.05, **: 0.001 < p < 0.01, ***: 1e-4 < p < 0.001).

### Overlapping of translated uORFs

Case studies using reporter assays have suggested that the CPuORFs in Homology Groups 1 and 3 can be regulated by their overlapping uORFs (Hanfrey et al., 2002; Wiese et al., 2004; Hanfrey et al., 2005). In our dataset, we detected thousands of transcripts that possess multiple translated uORFs (**Figure 9K**), with some uORFs overlapping with each other (e.g., **Figure 9L**). However, the extent of uORF overlapping at a genome-wide scale has remained unclear, especially since a comprehensive list of translated uORFs was previously lacking. As CiPS permits some levels of overlapping between uORFs, this offers an opportunity to identify uORF stacking globally. We systematically identified the overlapping events of translated uORFs on the most abundant transcript isoforms in our data, resulting in the identification of 681 uORFs involved in overlapping on 317 transcripts (**Dataset S6A**).

It is noteworthy that out of the 92 CPuORF transcripts detected in our data, 18 of them possess additional translated uORFs that stack with the CPuORF. Remarkably, we detected stacking uORFs in all five members of Homology Group 1 CPuORFs within *bZIP1*, *bZIP2*, *bZIP11*, *bZIP44*, and *bZIP53* mRNAs (**Dataset S6B**). This result correlates with the notion that the five members in Homology Group 1 contain an overlapping uORF upstream of the CPuORFs, which is also conserved (Wiese et al., 2004). Examples of uORF/CPuORF overlapping within *bZIP11* (Homology Group 1), *SAMDC2* (Homology Group 3), and *CIPK6* (Homology Group 27) are presented in **Figure 12A–C**. In *bZIP11*, three other uORFs stack with the CPuORF, but only the first uORF (18 aa) is translated at significant levels (**Figure 12A**). Similarly, in *SAMDC2*, one 3-aa uORF is translated and stacks with the CPuORF (**Figure 12B**), and in *CIPK6*, one 12-aa uORF is translated and stacks with the CPuORF (**Figure 12C**). These uORF stacking events, particularly the conserved stacking pattern in Homology Group 1, suggest that these upstream stacking uORFs could potentially regulate the translation of CPuORFs.

**Figure 12.**
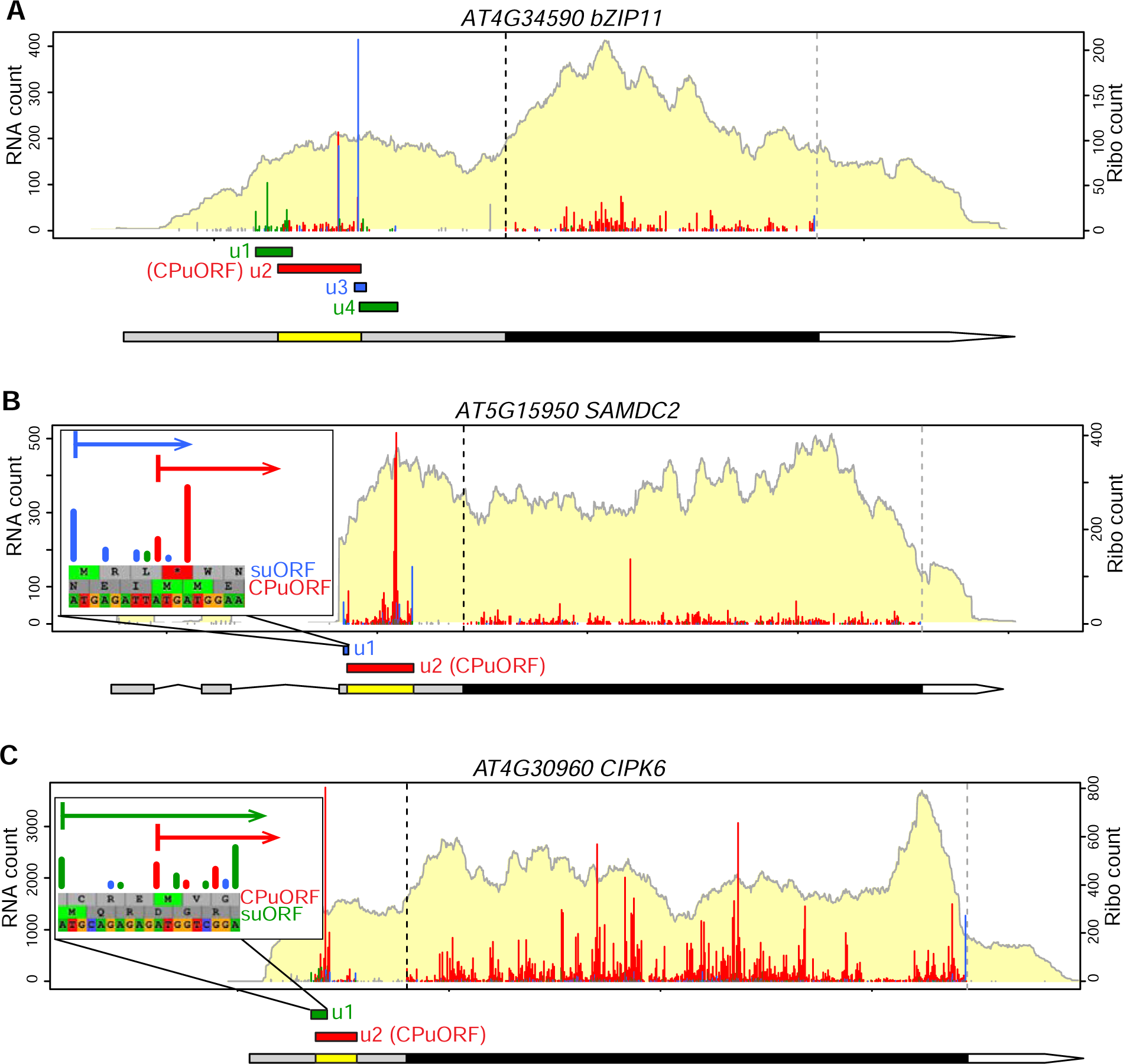
Overlapping of translated uORFs. (A–C) Translational profiles of three CPuORF genes show the overlapping of translated uORFs. The positions of CPuORFs (red boxes) and other uORFs (green or blue boxes, depending on their reading frame) are indicated. In (B–C), zoom-ins of the overlapping region between the CPuORF and the stacking uORF (suORF). Note *bZIP11* (A) is also shown in Figure 7B as an example of periodic ribosome stalling upstream of the CPuORF stop codon.

### Translated uORFs affected by alternative splicing

As the presence of potential uORFs is determined by the 5’ UTR sequences, the occurrence of translated uORFs could be regulated by alternative splicing in the 5’ UTRs. Two such examples have been reported in *HCS1*, a holocarboxylase synthetase, and *PIF3*, a repressor of light responses. In the case of *HCS1*, the presence or absence of the isoform-specific uORF leads to different translational initiations and alters the subcellular targeting of HCS1 (Puyaubert et al., 2008). In the case of *PIF3*, an intron retention within the 5’ UTR is regulated by Phytochrome B to repress the translation of the *PIF3* transcripts (Dong et al., 2020). Although the particular isoforms reported to affect the uORFs in *HCS1* and *PIF3* were not significantly expressed under our experimental conditions (**Figure S12A–B**), by examining the two most abundant isoforms of expressed genes in our data, we identified 594 translated uORFs that are affected by alternative splicing within 399 genes (**Dataset S7A, 7B**).

These uORFs are linked to specific isoforms through various alternative splicing events, including intron retention, alternative 5′ donor site, alternative 3′ acceptor site, and exon skipping (**Figure 13A–D**). Notably, *TOPLESS*, an important regulator in the auxin and jasmonic acid (JA) signaling, contains a cassette exon, which includes a translated uORF in isoform 2 (**Figure 13D**). These examples illustrate that the uORF presence and regulation could be widely controlled by alternative splicing and suggest the existence of isoform-specific translational control.

**Figure 13.**
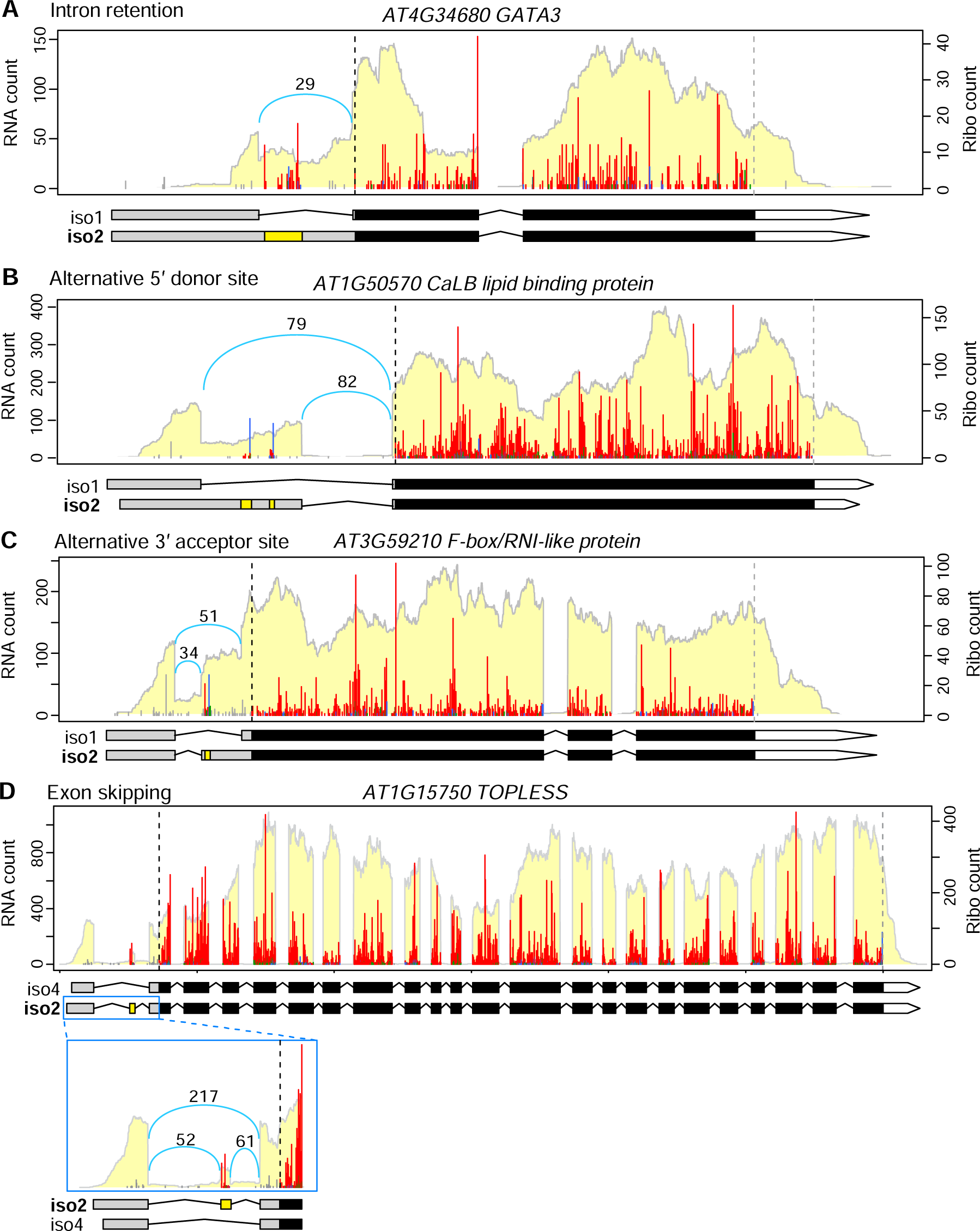
Translated uORFs regulated by alternative splicing. (A–D) Examples of translated uORFs affected by various types of alternative splicing. The number above the blue curly line indicates the read count across exon-exon junctions. The specific isoform number being plotted is indicated to the left of the gene model and bolded. The yellow boxes within the gene models represent the uORFs.

### Pathways regulated by uORFs

A prior bioinformatic analysis indicated that predicted uORFs are enriched in genes encoding transcription factors and protein kinases (Kim et al., 2007). To investigate what molecular functions and pathways are controlled by translated uORFs, we performed a GO term analysis on genes containing uORFs identified by RiboTaper or CiPS (**Figure S13**). Consistent with the previous prediction, we identified protein phosphorylation/modification, signal transduction, regulation of gene expression, etc. in the ‘Biological Process’ terms, and protein kinase activity, transcription regulator activity, DNA binding etc. in the ‘Molecular Function’ terms. Interestingly, many membrane-associated terms, including peroxisomal membrane, endosomes, Golgi, vesicles, plasma membrane etc. were identified in the ‘Cellular Component’ terms (**Figure S13**).

We also surveyed whether genes in various critical pathways harbor translated uORFs. Our analysis revealed that many key components in the light signaling, circadian clock, eight phytohormone signaling pathways (auxin, ABA, JA, ethylene, GA, SA, cytokinin, and brassinosteroid), development, as well as processes related to translational control, RNA biology, or signaling, contain translated uORFs (**Table 1**). These findings suggest that these important plant pathways are under translational control through uORFs. Therefore, our comprehensive uORF list serves as a valuable resource for further understanding gene expression regulation and plant growth.

**Table 1.**
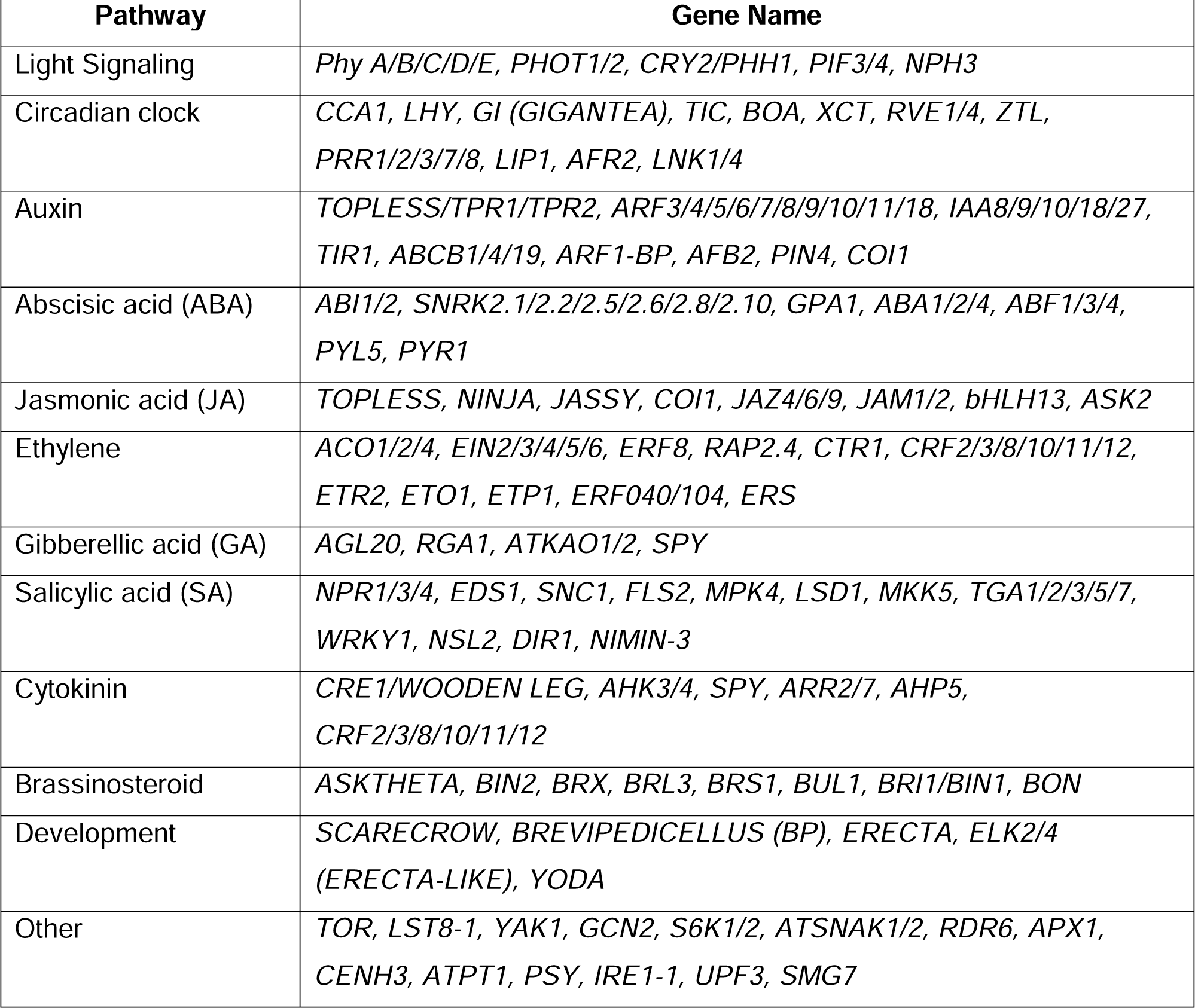
Important genes possessing translated uORFs.

## DISCUSSIONS

In this study, we tackled the identification of short ORFs by improving the coverage of Ribo-seq and developing a new computational pipeline for uORF identification. Our results provide a clearer view of the translational landscape and useful resources for diverse gene regulation studies, especially for uORFs on protein-coding transcripts and sORFs within presumed non-coding RNAs.

While a large portion of sORFs could be simply misannotated due to constraints of computational annotations, the translation of sORFs within well-characterized non-coding RNAs, such as primary transcripts of tasiRNAs and miRNAs, highlights the existence of ‘dual function’ RNAs (reviewed in (Ulveling et al., 2011; Raina et al., 2018)). That is, some RNAs may function as regulatory RNAs while possessing protein-coding capacity. In the case of tasiRNAs, the translation of the sORFs within the primary transcripts regulates the biosynthesis of tasiRNAs (Zhang et al., 2012; Yoshikawa et al., 2016; Li et al., 2016; Hou et al., 2016; Bazin et al., 2017; Iwakawa et al., 2021). In the case of miRNAs, application of synthetic micropeptides encoded by the sORFs in several pri-miRNAs increases the expression of the pri-miRNAs or mature miRNAs (Lauressergues et al., 2015; Sharma et al., 2020). For both tasiRNAs and pri-miRNAs, the translation and the regulatory RNA act in the same pathways, but it remains possible that the two functions act in different pathways, as shown in bacteria (reviewed in (Raina et al., 2018)). Only a limited number of sORFs have been characterized in plants so far, but they clearly have wide ranges of functions (Hanada et al., 2013; Tavormina et al., 2015; Hsu and Benfey, 2018; Fesenko et al., 2019; Ong et al., 2022). Studies in bacteria and mammals revealed that dozens of sORFs encode membrane proteins (reviewed in (Orr et al., 2020)). These small proteins could be a component of a large protein complex or involved in the recruitment of other proteins to membranes, the assembly of protein complexes within or near membranes, and controlling protein stability or activity of other proteins. The prediction that a significant portion of Arabidopsis sORFs we detected are secreted proteins points a promising direction for functional characterization of these sORFs.

We found that the out-of-frame (frame 2) RPFs at the codon preceding the stop interfere with the discovery of short uORFs. In eukaryotes, translational termination occurs when the ribosomal A-site encounters one of the three stop codons, followed by eRF1/eRF3 complex binding to the A-site and releasing the nascent peptide (reviewed in (Schuller and Green, 2018)). Cryogenic electron microscopy (Cryo-EM) analysis of mammalian ribosomes at termination revealed that eRF1 binding ‘pulls’ the nucleotide downstream of the stop codon into the A-site, thus packing 4 nts of mRNA at the A-site (Brown et al., 2015; Matheisl et al., 2015). We reasoned that the frame 2 RPFs at the codon preceding the stop are resulted from the structural rearrangement of ribosomes upon eRFs binding to the A-site during termination. Comparing the same length of RPFs from an elongating ribosome and a terminating ribosome, the packing caused by eRF1 binding would cause the RPF of a terminating ribosome to be 1 nt shorter at 5′ and 1 nt longer at 3′. This 1-nt shifting toward the 3′ causes the RPFs of terminating ribosomes to be mapped to the next reading frame (frame 2). Therefore, the frame 2 mapping of terminating RPFs we observed in Arabidopsis and tomato suggests that plant ribosomes behave like mammalian ribosomes at translation termination. Although plastid RPFs behave similarly to nuclear RPFs in multiple ways (**Figure 1A**, and **3A–B**), the frame 2 mapping at termination is lacking in plastid RPFs (**Figure 3B**). Currently, little is known about the mechanism of translation termination in plastid and mitochondria ribosomes, which have prokaryotic origins (Zoschke and Bock, 2018). In eukaryotes, eRF1 recognizes all three stop codons, whereas in prokaryotes, two different release factors recognize specific stop codons. Moreover, the release factors in eukaryotes and prokaryotes are evolutionarily unrelated (reviewed in (Buskirk and Green, 2017)). Our observations that plastid RPFs lack the frame 2 mapping at termination suggest that the mRNA packing caused by release factor binding may not occur in plastids. This phenomenon is likely the case for plant mitochondria as well, although specialized Ribo-seq optimizing for mitochondria ribosomes will be necessary to draw a clear conclusion.

Our CiPS pipeline, which accepts the frame 2 mapping of terminating RPFs as in frame, facilitates the identification of uORFs with relatively short lengths. Minimum uORFs in particular have only one codon and are therefore impossible to be found based on 3-nt periodicity. Given the codon usage of uORFs are similar to random triplet sequences in 5′ UTRs and most uORF peptides are not evolutionarily conserved, it is believed that most uORFs are selected for their regulatory role, rather than the encoded peptides (Von Arnim et al., 2014; Fields et al., 2015; Johnstone et al., 2016; Van Der Horst et al., 2020). Modulating uORFs has emerged as a promising approach to controlling gene expression and selection for agricultural traits (Xu et al., 2017; Zhang et al., 2018; Xing et al., 2020; Gage et al., 2022). As our results show that short uORFs are equally powerful as longer uORFs and most short uORFs are largely unexplored in previous studies, our uORF list offers new opportunities for gene regulation in diverse plant pathways, especially for genes encoding transcription factors and protein kinases, critical regulators in cellular signaling.

## MATERIALS AND METHODS

### Plant growth conditions and lysate preparation

Arabidopsis Col-0 seeds were surface sterilized with 70% ethanol for 5 min, followed by 33% bleach with 0.03% Tween 20 for 10 min, then rinsed with sterile water 5 times. The seeds were imbibed at 4°C in the dark for 2 days, then grown hydroponically in sterile liquid media (2.15 g/L Murashige and Skoog salt, 1% sucrose, 0.5 g/L MES, pH 5.7) while shaking at 85 rpm under a 16-h light (75–80 μmol m^−2^·s^−1^ from cool white fluorescent bulbs) and 8-h dark cycle at 22°C for 7 days. At Zeitgeber time 4 (4 hours after lights on), DMSO corresponding to 0.1% of the media volume was added to the media (these were control samples of our large-scale chemical treatment experiment). After 20 and 60 min, three biological replicates (∼300 plants per sample) were harvested at each time point and immediately flash frozen with liquid nitrogen.

Plant lysates were prepared as previously described (Hsu et al., 2016). Briefly, per 0.1 g of ground tissue powder was resuspended in 400 µL of lysis buffer (100 mM Tris-HCl [pH 8], 40 mM KCl, 20 mM MgCl_2_, 2% [v/v] polyoxyethylene [10] tridecyl ether [Sigma, P2393], 1% [w/v] sodium deoxycholate [Sigma, D6750], 1 mM dithiothreitol, 100 µg/mL cycloheximide [Sigma, C4859], 100 µg/mL chloramphenicol [Sigma R4408], and 10 units/mL DNase I [Epicenter, D9905K]). The lysates were spun at 3,000 g for 3 min, and the supernatant was transferred to a new tube and subsequently centrifuged at 20,000 g for 10 min. The supernatant was transferred to a new tube and the RNA concentration was determined with 10x dilutions using the Qubit RNA HS assay (Thermo Fisher Scientific, Q32852). Aliquots of 100 µL and 200 µL of the lysates were made, and they were flash frozen in liquid nitrogen and stored at −80°C until further processing.

### Ribo-seq library construction

Briefly, ribosome footprints were processed using 200 µL of the lysates described above, and sequencing libraries were constructed according to our previous method (Hsu et al., 2016) with a few modifications described below. Note the TruSeq Mammalian Ribo Profile Kit (illumina, RPHMR12126) and RiboZero Plant Leaf kit (Illumina, MRZPL1224) described here have been discontinued, researchers who are interested in our method could reference our custom library construction protocol (Wu and Hsu, 2022) instead.

The 200 µL of the lysates described above were treated with RNase I (50 units nuclease per 40 µg of RNA; the nuclease was included in TruSeq Mammalian Ribo Profile Kit, Illumina, RPHMR12126) for 1 hour at room temperature with gentle mixing. Then, 15 µl of SUPERase-IN (Invitrogen, AM2696) were added, and the lysate were passed through a size exclusion column (Illustra MicroSpin S-400 HR Columns; GE Healthcare, 27-5140-01). RNA > 17 nt was isolated with the RNA Clean & Concentrator-25 kit (Zymo Research, R1017) and separated on 15% urea-TBE gels (Invitrogen, EC68852BOX). Gel slices roughly between 27 and 31 nt were isolated, and the RNAs were purified as previously described. Next, rRNA depletion was performed using RiboZero Plant Leaf kit (Illumina, MRZPL1224) in one quarter of the recommended reaction volume. Ribo-seq libraries were constructed using the TruSeq Mammalian Ribo Profile Kit (illumina, RPHMR12126) as previously described with 9 cycles of PCR amplification. Libraries with equal molarity were pooled and sequenced on a Hi-Seq 4000 sequencer using single-end 50-bp sequencing.

The two major changes we made here were 1) reducing the RNA purification to one round before the size selection, and 2) changing the order of the rRNA depletion and size selection. In our previous protocol, we purified the RNA twice (first >17 nt and then <200 nt) to concentrate ribosome footprints before the rRNA depletion and gel purification. We reasoned that some footprints might be lost during the two rounds of RNA purification. In addition, the rRNA depletion by RiboZero limited the amount of input RNA that could be used according to the manufacturer’s recommendations. Therefore, we reasoned these two major changes could maximize the input and minimize the footprint loss during the processes.

### RNA-seq library construction

Total RNA greater than 200 nt was purified from 100 µL of the lysates described above using the RNA Clean & Concentrator-25 kit (Zymo Research, R1017) as previously described (Hsu et al., 2016). RNA integrity was evaluated using a Bioanalyzer (Agilent) RNA pico chip, and RNA integrity numbers (RINs) ranging from 7.2 to 7.7 were observed among the samples. A total of 4 µg of RNA per sample was subjected to rRNA depletion using the RiboZero Plant Leaf kit (Illumina, MRZPL1224) following the manufacturer’s recommendations. Then, 100 ng of rRNA-depleted RNA was fragmented to around 200-nt long based on the RIN reported by the Bioanalyzer, and strand-specific sequencing libraries were made using the NEBNext Ultra II Directional RNA Library Prep Kit (New England Biolabs, E7760S) with 8 cycles of amplification. Libraries of equal molarity were pooled and sequenced on a Hi-Seq 4000 using paired-end 100-bp sequencing.

### Sequencing data pre-processing and analysis

Data pre-processing and analysis were performed as previously described (Hsu et al., 2016), except that the Araport11 annotation (Cheng et al., 2017) was used in this study. Briefly, for Ribo-seq libraries, the adaptor (AGATCGGAAGAGCACACGTCT) was clipped with fastx_clipper (FASTX toolkit v0.0.14) (http://hannonlab.cshl.edu/fastx_tool-kit/). For both RNA-seq and Ribo-seq, we used Bowtie2 (v2.3.4.1) (Langmead and Salzberg, 2012) to remove rRNA, tRNA, snRNA, and snoRNA contaminant sequences.

### Transcriptome assembly, ORF identification using RiboTaper, statistical analysis and data visualization

For transcriptome assembly, the RNA-seq data from all six samples were first combined and mapped with STAR aligner (Dobin et al., 2013) with the following parameters -- alignIntronMax 5000, --alignIntronMin 15, --outFilterMismatchNmax 2, --outFilterMultimapNmax 2, --outFilterType BySJout, --alignSJoverhangMin 8 and --alignSJDBoverhangMin 2. The resulting bam file was used for reference-guided transcriptome assembly with Stringtie (Pertea et al., 2015) following our previous pipeline (Wu et al., 2019). We used gffcompare (Pertea and Pertea, 2020) to analyze Stringtie output gtf file and select the newly assembled transcripts (i.e., transcript types i, x, y, o, u and s).

Both the RNA-seq and Ribo-seq reads were mapped to the newly assembled gtf (**File S1**, which contains both Araport11 and newly assembled transcripts) with STAR aligner (RNA-seq parameters: identical to above mentioned prior to the transcriptome assembly; Ribo-seq parameters: --alignSJoverhangMin 4, --alignSJDBoverhangMin 1, --outSAMmultNmax 1, and the rest of the parameters were identical to RNA-seq). Combining all six samples, the total reads mapped to the genome for Ribo-seq and RNA-seq were 298.01 million and 180.67 million pairs, respectively.

For identifying translated ORFs, we used the bam files mapped above from both RNA-seq and Ribo-seq as input for RiboTaper (v1.3.1a) (Calviello et al., 2016). The Ribo-seq metaplot and the distribution of Ribo-seq reads in different genome features were generated using Ribo-seQC (Calviello et al., 2019) with 10% randomly selected reads. The Ribo-seq read lengths and offsets used in RiboTaper analysis were 24, 25, 26, 27, 28, and 8, 9, 10, 11, 12, respectively, as determined from the metaplots for nucleus- and chloroplast-encoded genes. The resulting unannotated ORFs and annotated mORFs were extracted from the RiboTaper output ORF_max_filt file. Since the P-site offsets for mitochondria-encoded genes are different from the nucleus- and chloroplast-encoded genes, the translational profiles of mitochondria-encoded genes were manually visualized using RiboPlotR.

All data visualization and statistical analysis were performed in R (v4.0.3) (Da Rosa et al., 2004). For Ribo-seq and RNA-seq data visualization, the P_sites_all files from RiboTaper were first processed using the following code: cut -f 1,3,6 P_sites_all | sort | uniq -c | sed -r ’s/^(*[^]+) +/\1\t/’ > output.txt to aggregate the read counts at each P-site. The Ribo-seq and RNA-seq profiles were plotted using RiboPlotR (Wu and Hsu, 2021). RiboPlotR presents the Ribo-seq data in the context of gene and transcript structure with exon-intron junctions, and the RPFs within ORFs are colorlZIcoded to indicate the reading frames.

### Proteomics sample preparation and analysis

Proteomic experiments were carried out using 4- and 21-day-old shoot and root as well as 12-day-old root tissue from Arabidopsis Col-0. The protein extraction and digestion using trypsin and Lys-C was carried out based on established methods (Song et al., 2018b, 2018a). Samples for datasets that include tandem mass tag (TMT) labeling and/or phosphopeptide enrichment were prepared as previously described (Clark et al., 2021; Montes et al., 2022; Song et al., 2020). Two-dimensional HPLC fractionation was performed using either strong cation exchange or basic reversed phase fractionation. Fractionated peptides were delivered to a Q Exactive Plus mass spectrometer using either an Agilent 1260 quaternary or a Thermo U3000 HPLC. Data dependent acquisition was performed using Xcalibur 4.0 software in positive ion mode with a capillary temperature of 275 °C and an RF of 60. MS1 spectra were measured at a resolution of 70,000 while MS2 spectra were measured at a resolution of 17,500 (label free) or 35,000 (TMT labeled). All raw data were analyzed together using MaxQuant version 1.6.7.0 (Tyanova et al., 2016). Spectra were searched against a custom protein database generated from the RiboTaper output file ORFs_max_filt (Dataset S1), which was complemented with reverse decoy sequences and common contaminants by MaxQuant. Carbamidomethyl cysteine was set as a fixed modification while methionine oxidation and protein N-terminal acetylation were set as variable modifications. Phospho (STY) was also set as a variable modification for samples that were phosphopeptide enriched. Digestion parameters were set to “specific” and “Trypsin/P;LysC”. For the TMT experiments the sample type was set to “Reporter Ion MS2”. Up to two missed cleavages were allowed. A false discovery rate less than 0.01 at both the peptide spectral match and protein identification level was required. The “second peptide” option was used to identify co-fragmented peptides.

### Calculate mORF translation efficiency

We first used STAR to map the RNA-seq and Ribo-seq reads to the CDS of annotated protein-coding genes. The resulting bam files were used to quantify the transcripts per million (TPM) of each gene using RSEM (v1.3.1) (Li and Dewey, 2011). Then, the mORF translation efficiency was calculated by dividing the Ribo-seq TPM to the RNA-seq TPM.

### uORF identification using CiPS (<u>C</u>ount, in-frame <u>P</u>ercentage and <u>S</u>ite)

The processed RiboTaper P-site file (i.e., output.txt, see above) was further analyzed for small uORFs identification. We first classified the blue signal at the (−1) codon as in frame. We then applied the following criteria to consider if a uORF is translated: 1) ≥ 10 RPF counts, 2) in-frame RPF percentage ≥ 50 % (identical to the RiboTaper cut-off), 3) in-frame RPF occupied site ≥ 30 % (allowing some tolerance if a uORF overlaps with other uORFs). For criterium 3, only the Ribo-seq occupied P-sites were evaluated. These cut-offs were empirically determined based on our data. For simplicity, for each gene, the two most abundant isoforms (determined by Kallisto (Bray et al., 2016)) with identical mORF starts were used for CiPS analysis. If the two isoforms had different mORF starts, the one with the more upstream mORF start was used for the analysis to avoid false positives. The duplicated uORFs shared by the two isoforms were removed.

To identify translated uORFs that overlap with each other, we used CiPS-identified uORFs within the most abundant isoforms for the analysis.

For uORFs regulated by alternative splicing, we considered the two most abundant isoforms and identified translated uORFs (detected by CiPS) that only exist in one of the two isoforms.

### Constructs for dual luciferase assays

For the dual luciferase plasmid, the 35S-promoter sequence upstream of *Renilla Luciferase* (*REN*) within the pGreen II 0800 Luc plasmid (Hellens et al., 2005) was synthesized by BioBasic and cloned into the upstream of the *Firefly Luciferase* (*FLUC*) via Kpn I and Xho I. The resulting construct was renamed pHsu-133. All 5′ UTR sequences tested were synthesized by BioBasic and cloned into pHsu-133 between the 35S-promoter and the *FLUC* via BamH I and Not I. The exact 5′ UTR sequences tested were listed in **File S2**. For *S6K1*, *RGA1* constructs, annotated 5′ UTR sequences containing the wild-type or mutated uORF (ATG was changed to AGG) were used. For minimum uORF constructs, annotated 5′ UTR sequences containing the wild-type or mutated uORF (ATG was changed to ATC) were used. For tiny uORF constructs, 5′ UTR sequences containing the wild-type or mutated uORF (ATG was changed to AGG), including 40 nt upstream of the uORF to the end of 5′ UTR, were used.

### Dual luciferase assays and qRT-PCR

Arabidopsis protoplast preparation and transformation were modified from (Reis et al., 2020). Protoplasts were isolated from 20 to 21-day-old Col-0 fully expanded rosette leaves grown on soil under a 16-h light (∼100 μmol m^−2^·s^−1^ from cool white fluorescent bulbs) and 8-h dark cycle at 22°C. Finely cut leaf slices were immersed in enzyme solution (1% [w/v] cellulase, 0.25% [w/v] macerozyme, 0.4 M mannitol, 20 mM KCl, 20 mM MES, and 10 mM CaCl_2_) followed by vacuum infiltration for 30 minutes and gentle shaking at 40 rpm in the dark for 2–2.5 hours to release the protoplasts. The protoplasts were passed through a 70 µM cell strainer, centrifuged at 100 g at 4°C for 3 minutes, and washed with cold W5 solution (154 mM NaCl, 125 mM CaCl_2_, 5mM KCl, and 2 mM MES) twice. The protoplasts were counted and resuspended in MMG solution (4.5 mM MES [pH 5.7], 0.4 M mannitol, and 15 mM MgCl_2_) at 10^5 protoplasts/150 µL. For protoplast transformation, 10^5 protoplasts were combined with 5 µg plasmid DNA and 170 µL polyethylene glycol (PEG) solution (40% [w/v] PEG4000, 0.2 M mannitol, and 100 mM CaCl_2_), and incubated for 5 minutes. After consecutive washing with W5 solution, the transformed protoplasts were incubated in the dark for 16 to 18 hours. Typically, 8 replicates of transformation were performed for each plasmid DNA in one experiment, and the experiments were repeated twice or three times with similar results. The plasmid DNA was prepared using ZymoPURE II Plasmid Midiprep Kit (Zymo #D4201). After overnight incubation, protoplasts were harvested by centrifugation at 2250 g for 3 minutes at 4°C. The lysates were generated by adding 100 µL 1X Passive Lysis Buffer included in the Dual-luciferases assay kit (Promega E1960) to the protoplasts and vigorously shaking at room temperature for 15 minutes. The lysates were cleared by centrifuging at 2250 g for 3 min, and 20 µL of the supernatant was used for the dual-luciferases assay measured in GloMax Navigator Plate Reader (Promega, GM2010) as specified by the manufacturer. The FLUC luminescence was normalized to their corresponding REN luminescence.

For quantifying *FLUC* and *REN* transcript levels, 8 µL 10% SDS was added to the remaining 80 µL lysates above, and total RNA was isolated using Zymo RNA Clean and Concentrator Kit (Zymo, R1014). RNA was converted to cDNA using LunaScript RT SuperMix (New England Biolabs, 3010) in a final 10 µL reaction. Ten-fold diluted cDNA was used in qRT-PCR using Luna Universal qPCR Master Mix (New England Biolabs #M3003E) in a final 10 µL reaction in QuantStudio 3 Real-Time PCR machine (Thermo Fisher Scientific). Primers for *FLUC* and *REN* were previously described in (Zhang et al., 2018). The quantification was determined using the standard curve method, and the relative *FLUC* transcript levels were normalized to their corresponding *REN* transcripts levels.

## AVAILABILITY OF DATA AND MATERIALS

All raw and processed sequencing data generated in this study have been submitted to the NCBI Gene Expression Omnibus (GEO; https://www.ncbi.nlm.nih.gov/geo) under accession number GSE183264. The original MS proteomics raw data may be downloaded from MassIVE (http://massive.ucsd.edu) using the identifier “MSV000085044”; the three key files (i.e., evidence.txt, peptides.txt and proteinGroups.txt) are located in the /search/combined/txt subfolder.

## AUTHOR CONTRIBUTIONS AND ACKNOWLEDGEMENTS

HLW and PYH designed the research and interpreted the data. PYH performed the sequencing experiments. HLW analyzed the sequencing data and prepare figures and tables for the paper. QA and RT performed dual luciferase assays. QA performed qRT-PCR. GS and JME generated the proteomics datasets. CM configured the mass-spec database and performed the search. PYH and JW supervised the research. HLW and PYH wrote the manuscript with input from all other authors.

We thank Phong Nguyen for his assistance in dual luciferase assays and qRT-PCR; we thank Rodrigo Reis and Yves Poirier for sharing their protocol for dual luciferase assays, and we thank Isaiah Kaufman for his critical review of this manuscript. pGreen II 0800 Luc plasmid is a gift from Roger Hellens. This work used the Vincent J. Coates Genomics Sequencing Laboratory at UC Berkeley, supported by an NIH S10 OD018174 Instrumentation Grant. This work was supported by a National Science Foundation grant (2051885) to PYH and Iowa State University Plant Science Institute, USDA NIFA Hatch project IOW3808, and National Science Foundation grant (IOS-1759023) to JWW.

