## Supplemental figures for "Improved Super-Resolution Ribosome Profiling Revealed Prevalent Translation of Upstream ORFs and Small ORFs in Arabidopsis"

**Figure S1** (supporting Figure 1)

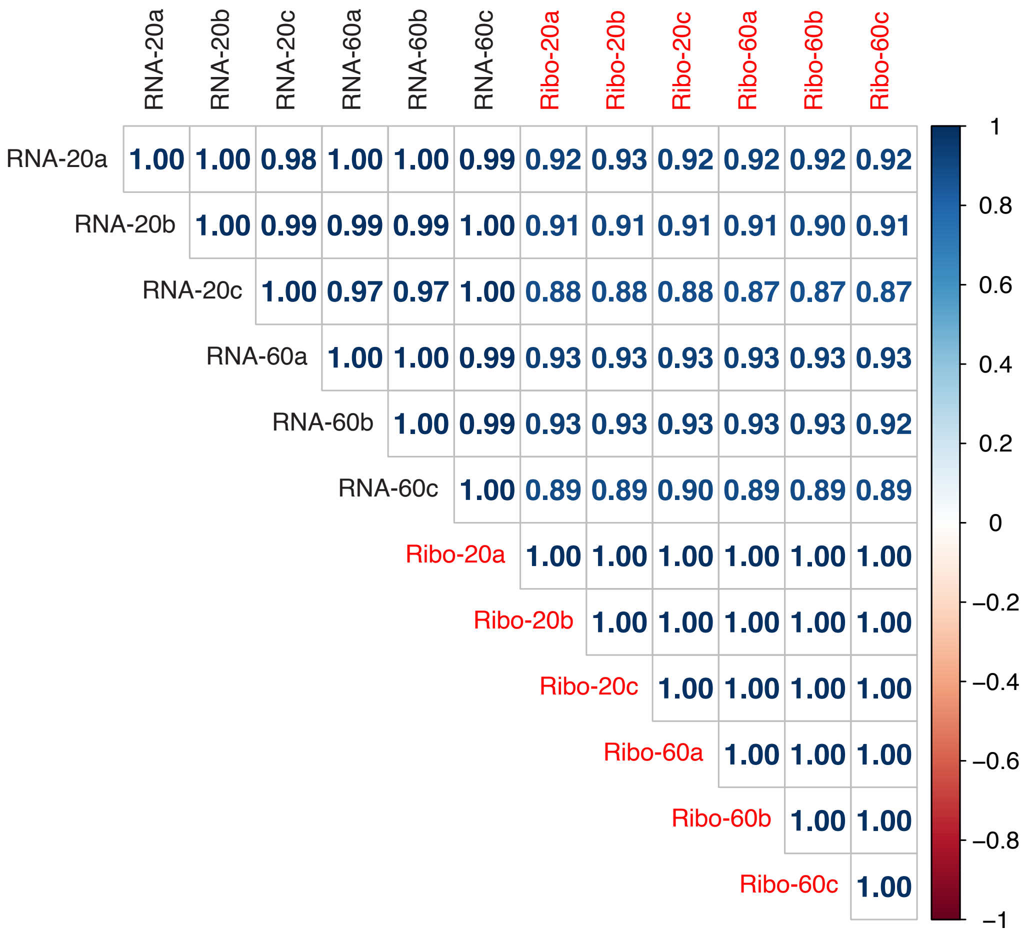


**Figure S1.** Pearson correlations among six samples. RNA-seq and Ribo-seq read counts within the annotated CDS of nucleus-encoded protein-coding genes are presented.

**Figure S2.** ORF identification workflow. (supporting Figure 2)

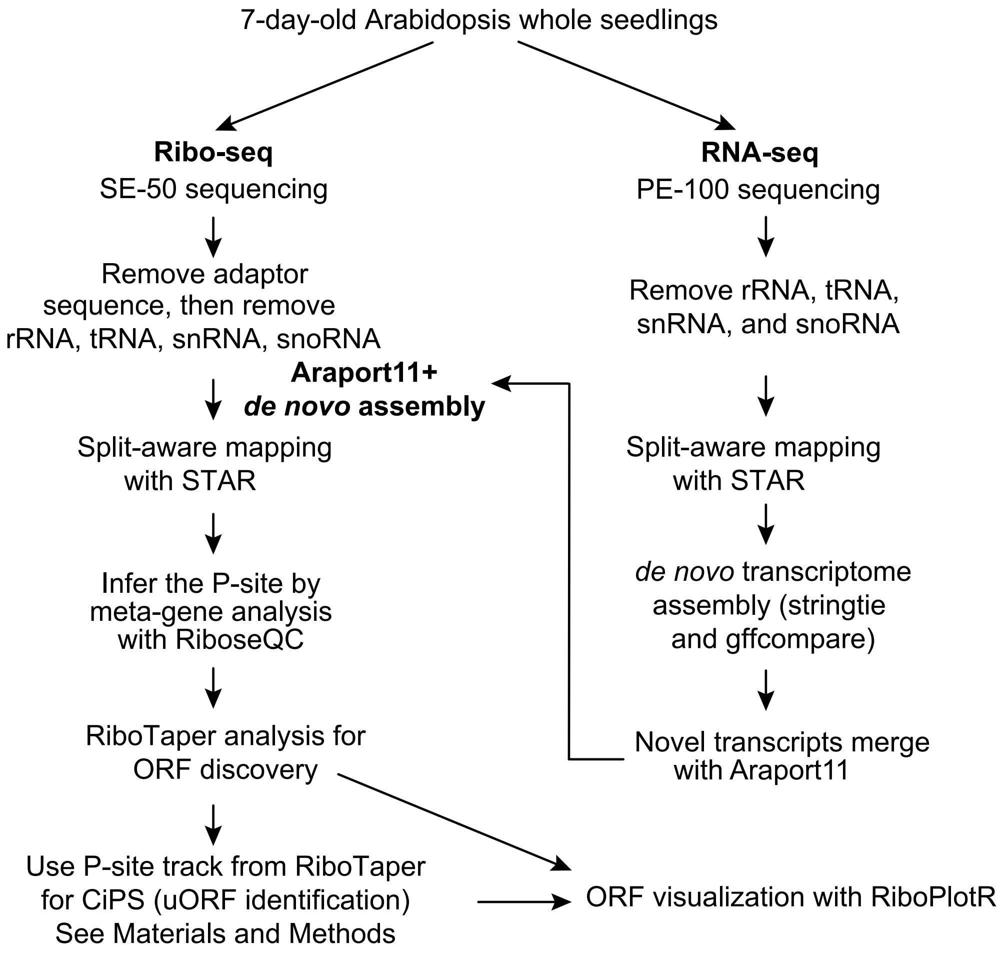

**Figure S3** (supporting Figure 2)

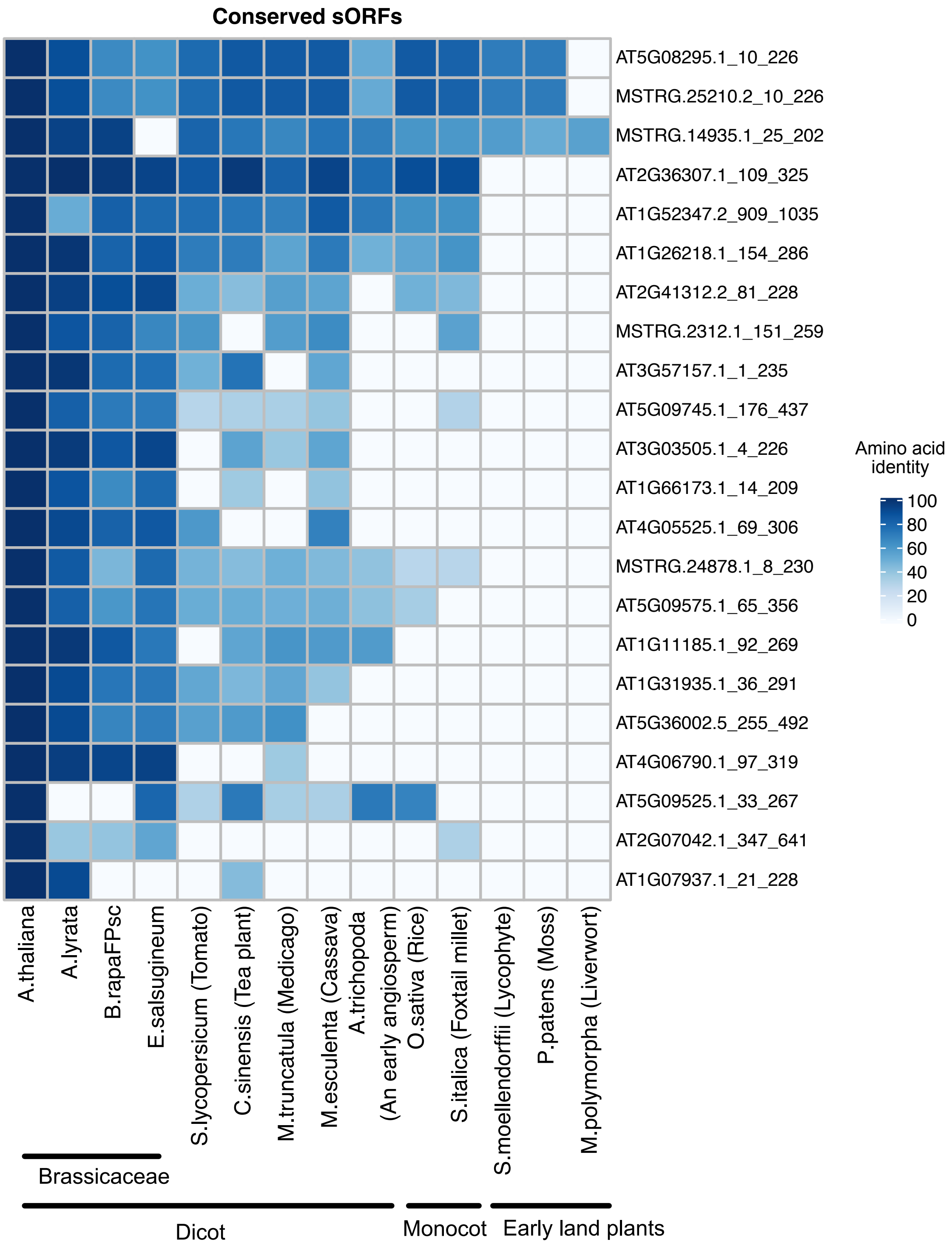


**Figure S3.** The evolutionary conservation of single-exon sORFs.

**Figure S4** (supporting Figure 3)

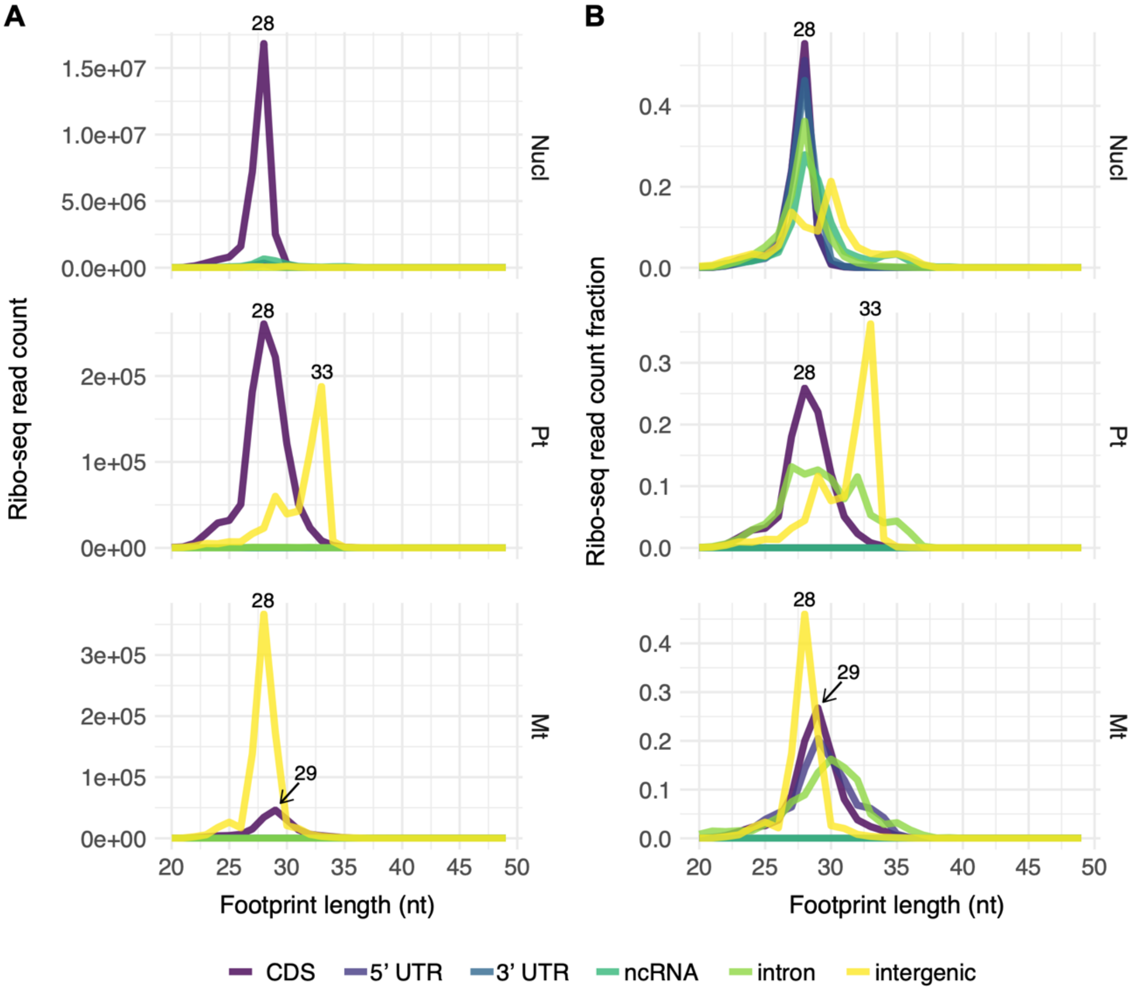


**Figure S4.** Ribosome footprint lengths and genomic features mapped to nucleus-, plastid-, and mitochondria-encoded genes. 10% of randomly selected reads were used for analysis. The output of Ribo-seQC (Calviello et al., 2019) is presented.

**Figure S5** (supporting Figure 3)

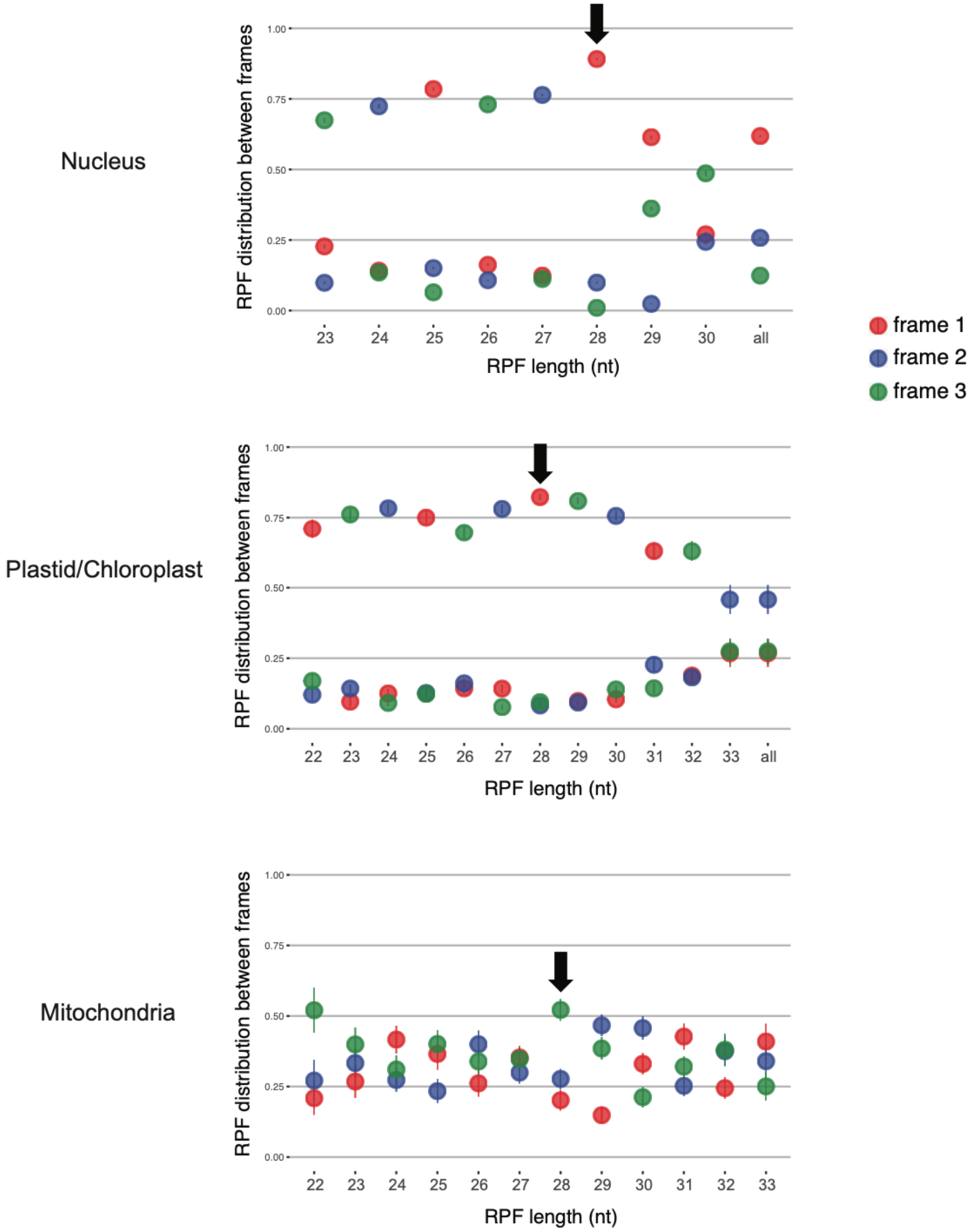


**Figure S5.** 3-nt periodicity observed in individual RPF lengths for nucleus-, plastid-, and mitochondria-encoded genes. The output of Ribo-seQC (Calviello et al., 2019) is presented. The RPF length exhibits the best 3-nt periodicity is highlighted by the black arrow.

**Figure S6** (supporting Figure 6)

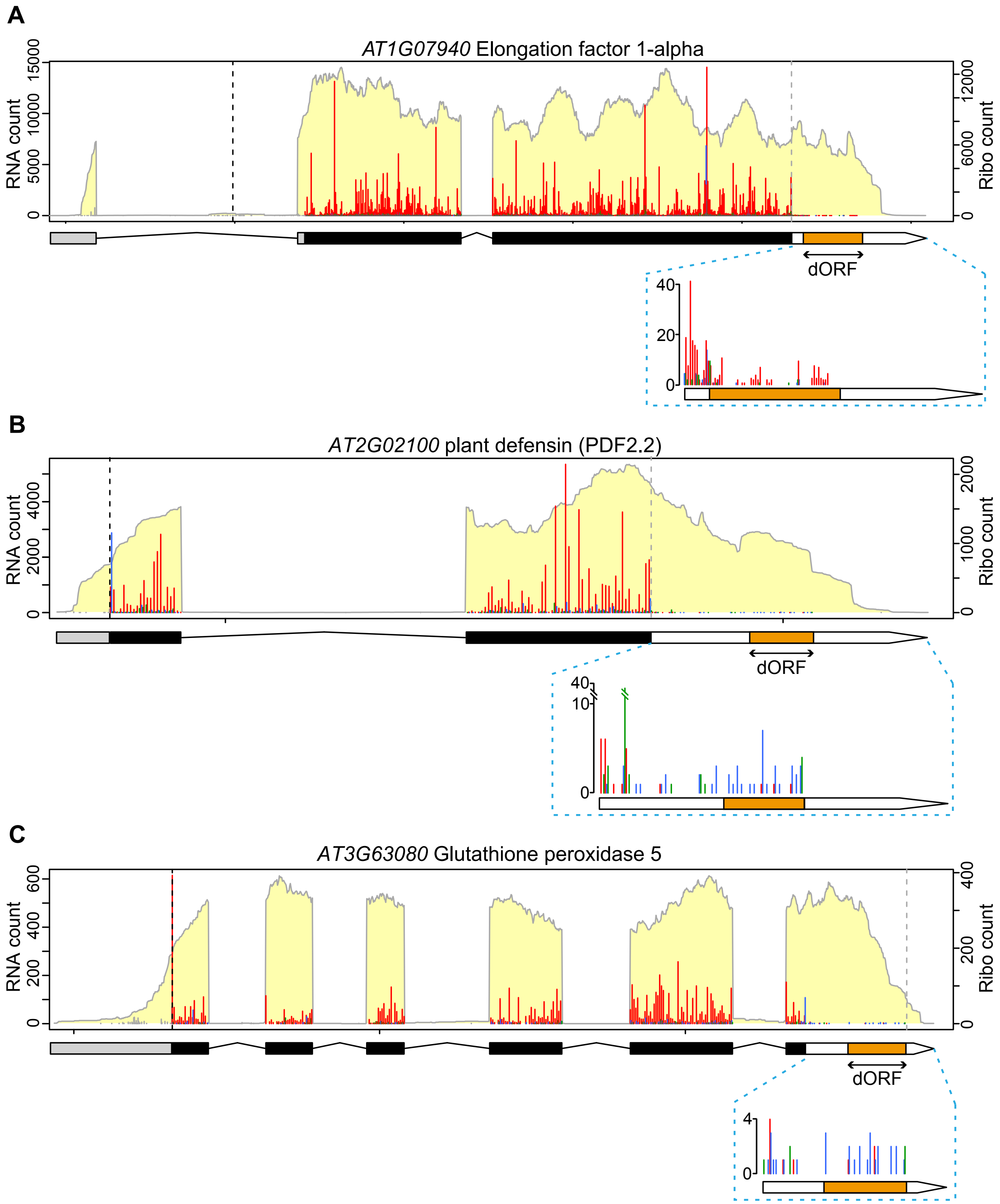


**Figure S6.** Zoom-ins of type 1 dORFs (Figure 6B–D) showing potential readthrough from the mORF (red reading frame) or an upstream alternative ORF (blue reading frame).

**Figure S7** (supporting Figure 6)

**A** *AT3G46640 LUX transcription factor* **(shown in Fig. 6E)**

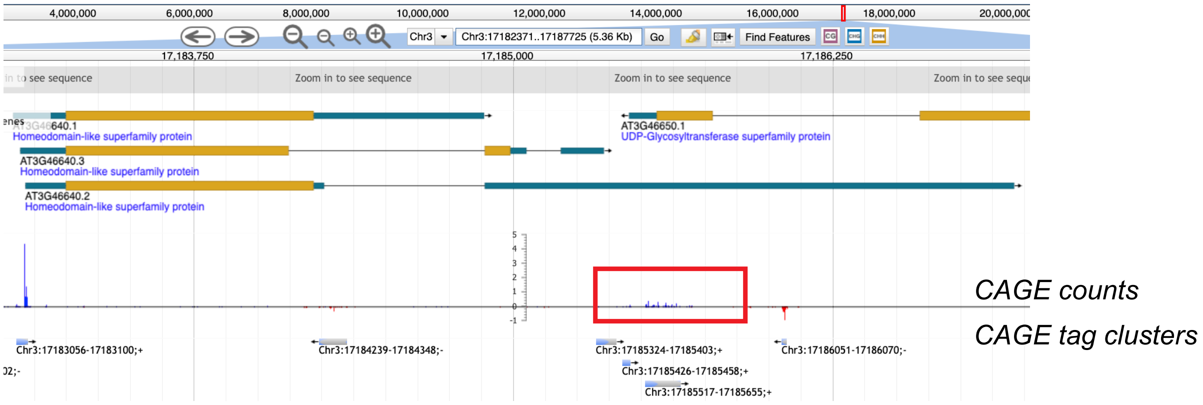


**B** *AT1G49980 DNA/RNA polymerases superfamily protein* **(shown in Fig. 6F)**

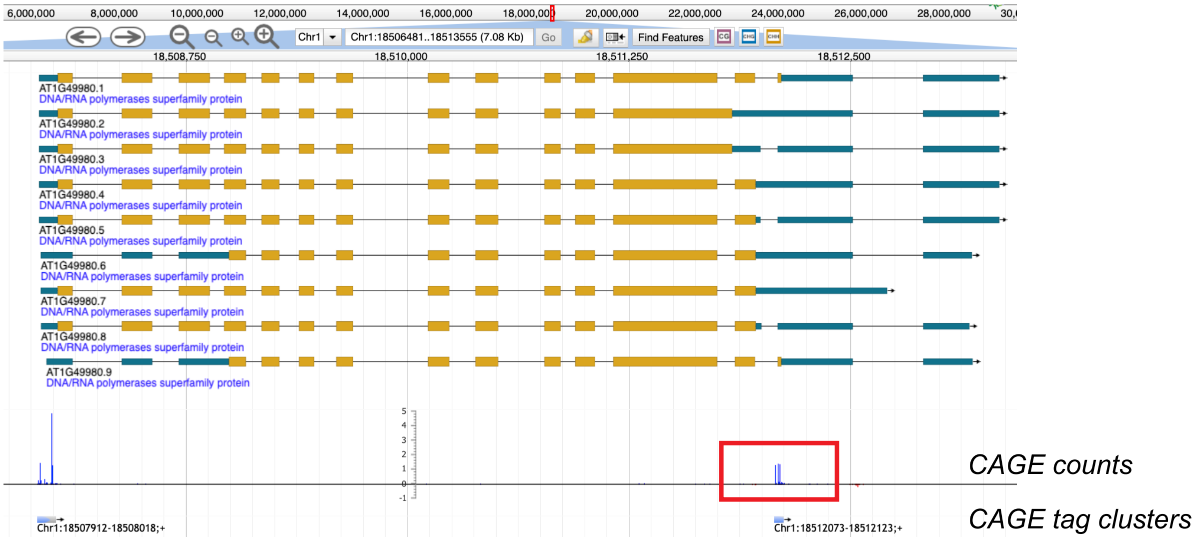


**Figure S7.** Type 2 dORFs have independent transcription start sites (highlighted by red boxes) supported by published CAGE data (Thieffry et al., 2020).

**Figure S8** (supporting Figure 6)

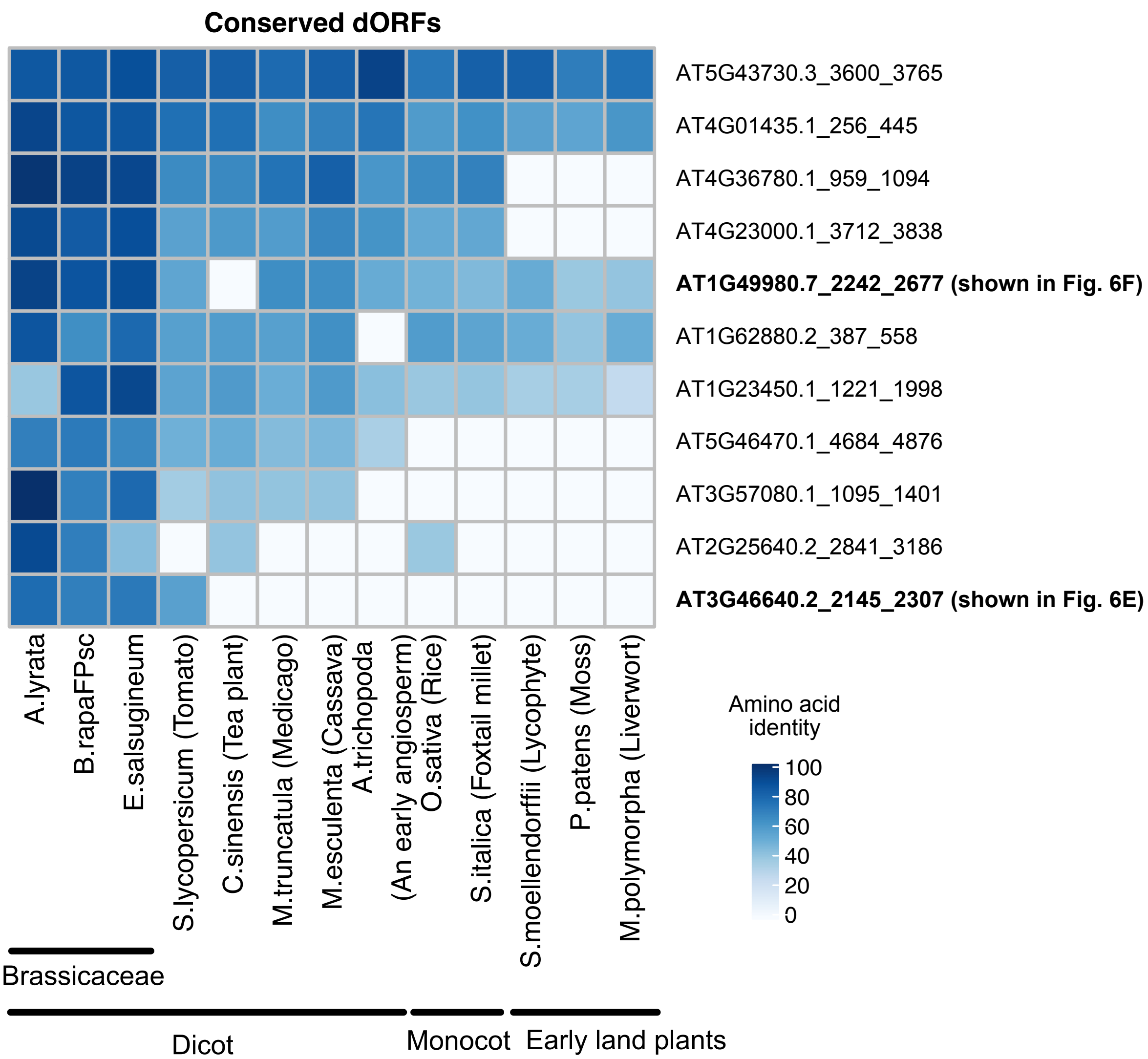


**Figure S8.** The evolutionary conservation of dORFs.
A tBLASTn search of the dORFs (≥ 20 aa) identified 11 of them have homologs outside of the Brassicaceae. These 11 dORFs all fall into type 2 dORFs, suggesting they were identified as dORFs by RiboTaper due to annotation issues.

**Figure S9** (supporting Figure 10)

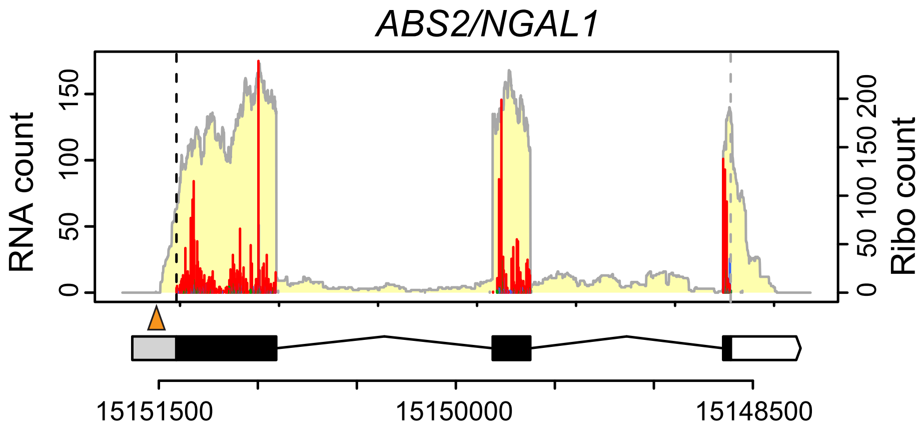


**Figure S9.** The minimum uORF reported in *ABS2* was not detected in our experimental conditions. *ABS2* uses a transcription start site downstream of the annotated transcription start and the minimum uORF (orange triangle).

**Figure S10** (supporting Figure 11)

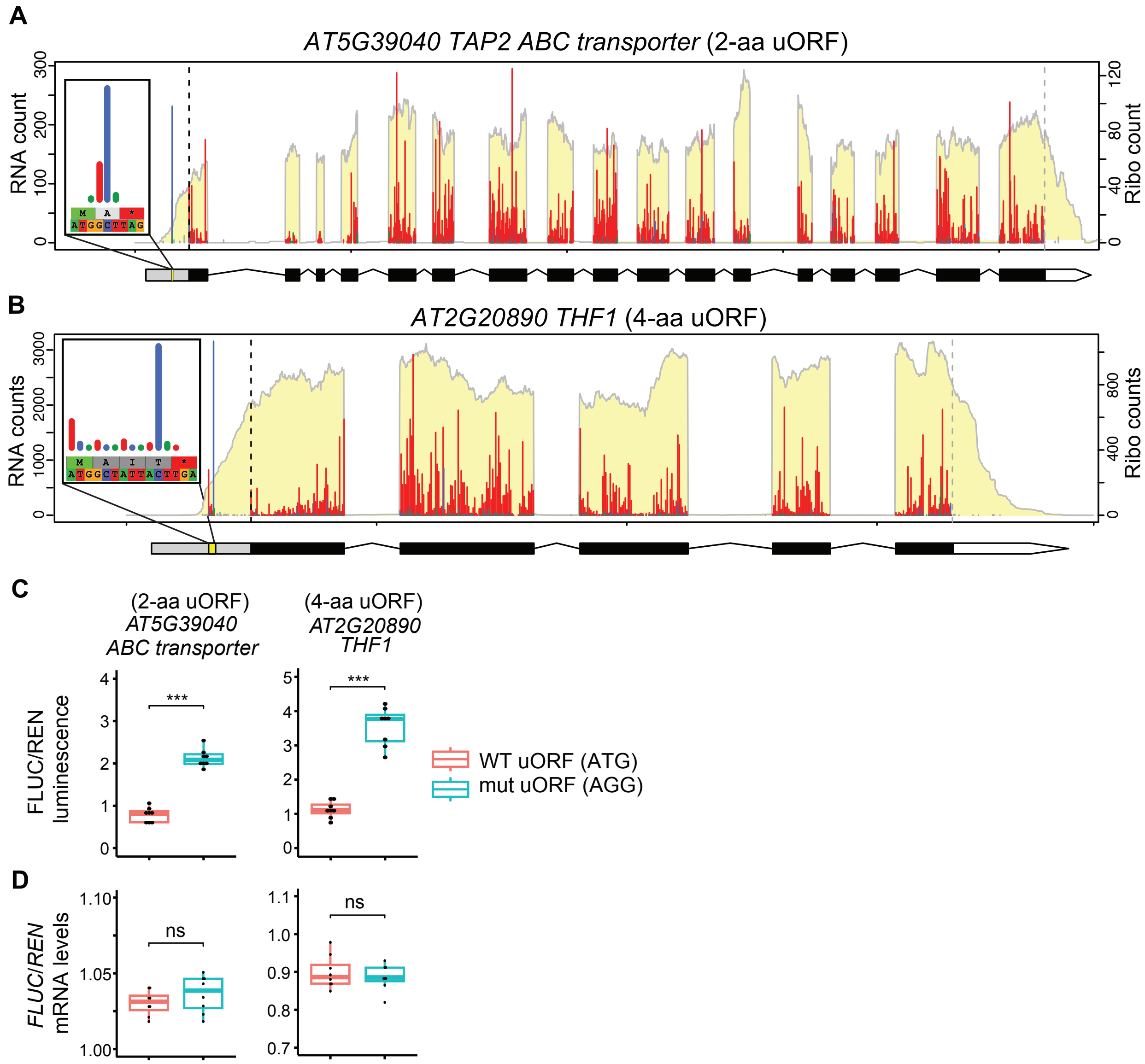


**Figure S10.** Additional examples of tiny uORFs and dual-luciferase assay.

(A–B) Examples of tiny uORFs of varying uORF length.

(C) Relative FLUC luminescence comparing 5′ UTRs carrying the wildtype uORF (ATG) or mutated uORF (AGG). FLUC luminescence levels are normalized to REN luminescence levels.

(D) Relative *FLUC* mRNA levels comparing 5′ UTRs carrying the wildtype (ATG) tiny uORF or mutated (AGG) tiny uORF. *FLUC* mRNA levels are normalized to *REN* mRNA levels. The statistical significance for boxplots in (C–D) was determined by Wilcoxon rank sum test (*: $0.01 < p < 0.05$, **: $0.001 < p < 0.01$, ***: $1e-4 < p < 0.001$).

**Figure S11** (supporting Figure 11)

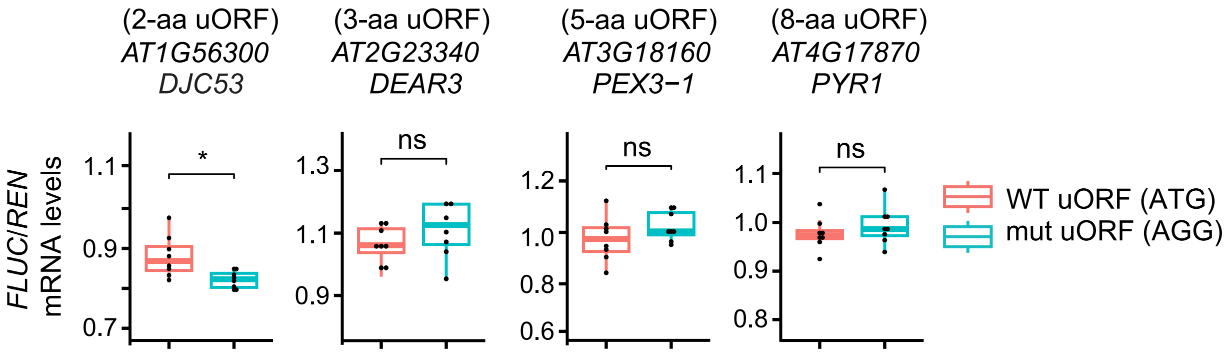


**Figure S11.** Relative *FLUC* mRNA levels comparing 5′ UTRs carrying the wildtype (ATG) tiny uORF or mutated (AGG) tiny uORF. *FLUC* mRNA levels are normalized to *REN* mRNA levels. The statistical significance was determined by Wilcoxon rank sum test (*: $0.01 < p < 0.05$, **: $0.001 < p < 0.01$, ***: $1e\text{-}4 < p < 0.001$).

**Figure S12** (supporting Figure 12)

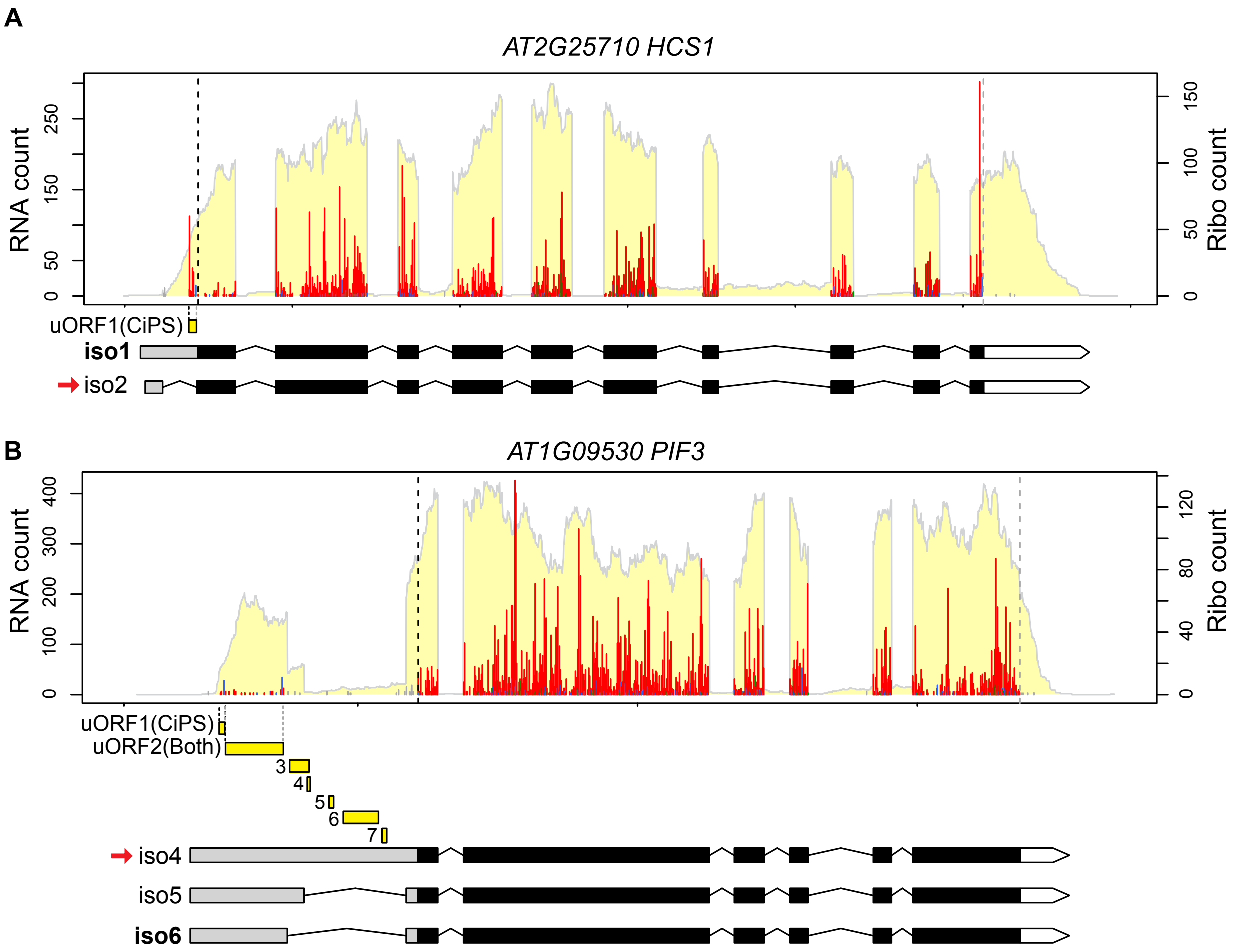


**Figure S12.** The specific isoform affecting uORFs in *HCS1* and *PIF3* (Puyaubert et al., 2008; Dong et al., 2020) are not significantly expressed in our experimental conditions. Previously reported isoforms affecting uORF presence are marked by a red arrow next to the gene models. Translated uORFs identified by CiPS-only, or by both RiboTaper and CiPS, as well as potential uORFs, are indicated.

**Figure S13** (supporting Table 1)

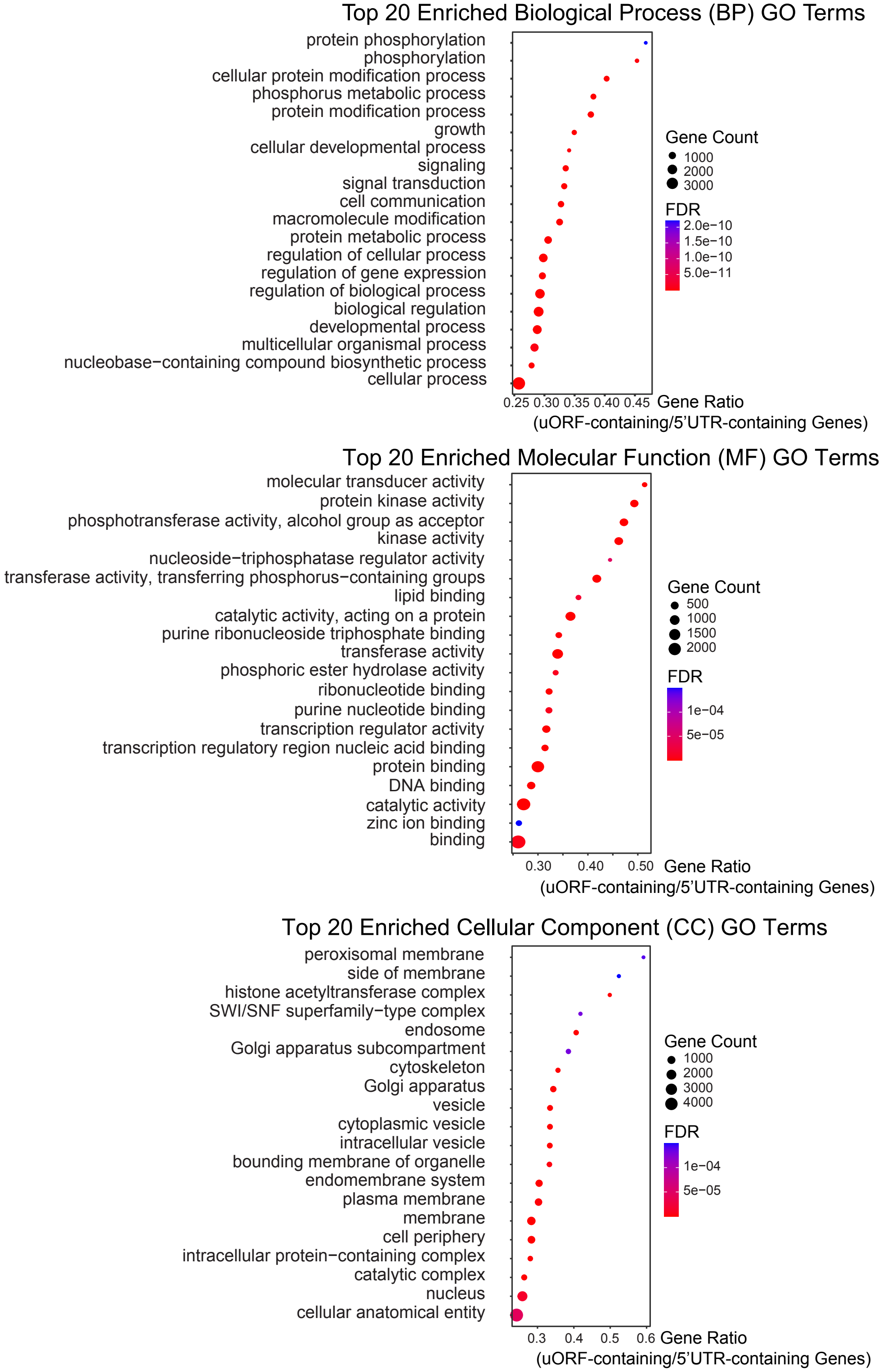


**Figure S13.** GO-term analysis of translated uORF genes identified by RiboTaper or CiPS. Top 20 enriched GO terms in Biological Process, Molecular Function, and Cellular Component are presented. Gene count is shown by circle size and FDR is shown by various colors as indicated.
